# Polygenic Risk for Alcohol Use Disorder Affects Cellular Responses to Ethanol Exposure in a Human Microglial Cell Model

**DOI:** 10.1101/2024.02.19.581066

**Authors:** Xindi Li, Jiayi Liu, Andrew J. Boreland, Sneha Kapadia, Siwei Zhang, Alessandro C. Stillitano, Yara Abbo, Lorraine Clark, Dongbing Lai, Yunlong Liu, Peter B Barr, Jacquelyn L. Meyers, Chella Kamarajan, Weipeng Kuang, Arpana Agrawal, Paul A. Slesinger, Danielle Dick, Jessica Salvatore, Jay Tischfield, Jubao Duan, Howard J. Edenberg, Anat Kreimer, Ronald P. Hart, Zhiping P. Pang

**Author notes:** **Correspondence** and requests for materials should be addressed to Zhiping P. Pang.

## Abstract

Polygenic risk scores (PRS) assess genetic susceptibility to Alcohol Use Disorder (AUD), yet their molecular implications remain underexplored. Neuroimmune interactions, particularly in microglia, are recognized as significant contributors to AUD pathophysiology. We investigated the interplay between AUD PRS and ethanol in human microglia derived from iPSCs from individuals with high- or low-PRS (HPRS or LPRS) of AUD. Ethanol exposure induced elevated CD68 expression and morphological changes in microglia, with differential responses between HPRS and LPRS microglial cells. Transcriptomic analysis revealed expression differences in MHCII complex and phagocytosis-related genes following ethanol exposure; HPRS microglial cells displayed enhanced phagocytosis and increased *CLEC7A* expression, unlike LPRS microglial cells. Synapse numbers in co-cultures of induced neurons with microglia after alcohol exposure were lower in HRPS co-cultures, suggesting possible excess synapse pruning. This study provides insights into the intricate relationship between AUD PRS, ethanol, and microglial function, potentially influencing neuronal functions in developing AUD.

## Introduction

Alcohol Use Disorder (AUD) remains a substantial contributor to the global burden of disease, posing an elevated risk for premature mortality and disability (*1*). The etiology of AUD is multifaceted, with genetic factors accounting for approximately half of the inter-individual variation in susceptibility (*2*). This heritable dimension of AUD is highly polygenic (*3-6*). Polygenic risk scores (PRS) summarize the combined impact of numerous genetic variants on an individual’s risk for specific diseases. AUD PRS capture a fraction of the genetic risk (*7-12*), but how that might affect cellular functions is unknown.

Immune and inflammatory pathways in the brain are activated in AUD (*13*). In gene expression profiling of postmortem human brains, a connection between immune responses and AUD has been established; genes exhibiting significant expression differences are enriched in pathways related to interferon signaling (*14*). Interferons, with their antiviral, antiproliferative, and immunomodulatory effects, play a critical role in human innate and adaptive immune responses during chronic alcohol exposure (*14*). Using human induced pluripotent stem cells (iPSCs) as a model system, it has been shown that ethanol exposure activates NLRP3 inflammasome in human iPSCs and iPSC-derived neural progenitor cells (*15*). Microglia are the primary resident immune cells in the brain (*16*) and possess a variety of receptors that enable them to sense alterations in their microenvironment, leading to changes in transcription, morphology, and function (*17, 18*). Brain microglial cells play multifaceted roles, including the regulation of inflammation, participation in phagocytosis (*19*), and engagement in specialized brain-specific functions through interactions with neurons, such as the modulation of neurotransmission and synaptic pruning (*20, 21*). Exposure to ethanol triggers microglia activation, leading to morphological changes and heightened immune responses in rodent models (*22-26*). Microglial depletion has been shown to mitigate alcohol-dependence-associated behaviors, impair synaptic function, and reverse expression changes in alcohol-dependent inflammatory-related genes in mice (*27*). It has also been suggested that microglia-mediated synaptic pruning may constitute the underlying mechanism behind synapse loss and memory impairment induced by prolonged alcohol consumption (*28*).

It is increasingly evident that rodent microglia may not faithfully mirror the biology of their human counterparts, as recent transcriptomic studies have revealed substantial differences, including the abundant expression of specific immune genes in human microglia that are not part of the mouse microglial signature (*17, 29, 30*). Human microglia can be derived from human iPSCs (*30-34*); they serve as an excellent model system for studying neuroimmune interactions (*30, 35, 36*). When human iPSCs differentiated into microglia were exposed to brain substrates, including synaptosomes, myelin debris, apoptotic neurons, or synthetic amyloid-beta fibrils, it resulted in a variety of transcriptional changes (*37*). These changes corresponded to gene signatures found in human brain microglia, particularly those enriched in neurodegenerative diseases (*37*). Therefore, human iPSC-derived microglia could be used as an in vitro model to understand disease-relevant cellular phenotypes. Despite the involvement of microglia in the pathophysiology of AUD (*27, 38-40*), the specific cellular and molecular mechanisms that govern the interaction between ethanol exposure and AUD PRS in the human context remain unknown.

In this study, subject-specific microglia were generated from iPSC lines derived from selected participants with AUD high PRS (HPRS, n=8, top 75 percentile, 4 males, 4 females) or low PRS (LPRS, n=10, bottom 25 percentile, 5 males and 5 females). These were from the Collaborative Study on the Genetics of Alcoholism (COGA). COGA has found many genetic and molecular mechanisms underlying the functional alteration in AUD (*41*). We selected subjects with a defined clinical diagnosis of AUD with HPRS or control samples LPRS with no AUD(*12*), which provide a unique opportunity to investigate the gene and environment (i.e., ethanol exposure) interaction and cellular functions. Since PRS summarizes a large number of genomic variations and there are no consistent variants defining each group, we expect to find a large degree of variability among cell lines within each group. In highly enriched human microglial cells derived from iPSCs, we found that ethanol exposure leads to upregulation of CD68 expression and alterations in microglial morphology, with distinct effects observed in microglia derived from subjects with different AUD PRS. Transcriptomic analysis unveiled differences between HPRS and LPRS microglial cells in the expression of genes related to the MHCII complex and to phagocytosis after ethanol exposure. Furthermore, an enhanced phagocytic process with increased *CLEC7A* (C-type lectin domain family 7 member A) RNA expression was observed in HPRS microglial cells following ethanol exposure, which was not observed in LPRS microglial cells. Our study reveals the intricate interplay between AUD-related polygenetic factors and microglial function in AUD, offering novel insights into the mechanism underlying AUD in humans.

## Results

### Generation of human iPSC-derived microglia

To test the hypothesis that AUD PRS affects cellular mechanisms, we prepared 18 iPSC lines from lymphocytes and lymphoblastoid cells from participants collected by the Collaborative Studies on Genetics of Alcoholism (COGA) (*42-44*) based on their AUD PRS (*12*). Eight participants (4 men and 4 women) had AUD and a PRS > 75 percentile (HPRS, 7 of which were above 90 percentile), and 10 (5 men, 5 women) were unaffected participants with AUD < 25 percentile (LPRS, 9 below 10 percentile) (**Table S1**). The pluripotency of these iPSC lines was validated with immunohistochemical staining with Oct4 and SOX2 (**Fig.S1**).

Using an established, highly efficient, and reproducible protocol (*45*), we generated microglial cells from these iPSCs. We first derived primitive macrophage progenitors (PMPs) from iPSCs (**Fig. 1A, B, Table S1**). Across multiple iPSC lines, more than 90% of the PMPs exhibited expression of the hematopoietic progenitor cell markers CD235 (**Fig. 1C, D**) and CD43 (**Fig. 1E, F**). After confirming the PMP cell identities, we differentiated these PMPs into microglial cells by adding IL-34 and GM-CSF for one week. Microglial identity was confirmed through immunostaining for a panel of microglia-specific markers, including IBA1, P2Y12, CX3CR1, TMEM119, and PU.1 (**Fig. 1G**).

**Fig. 1.**
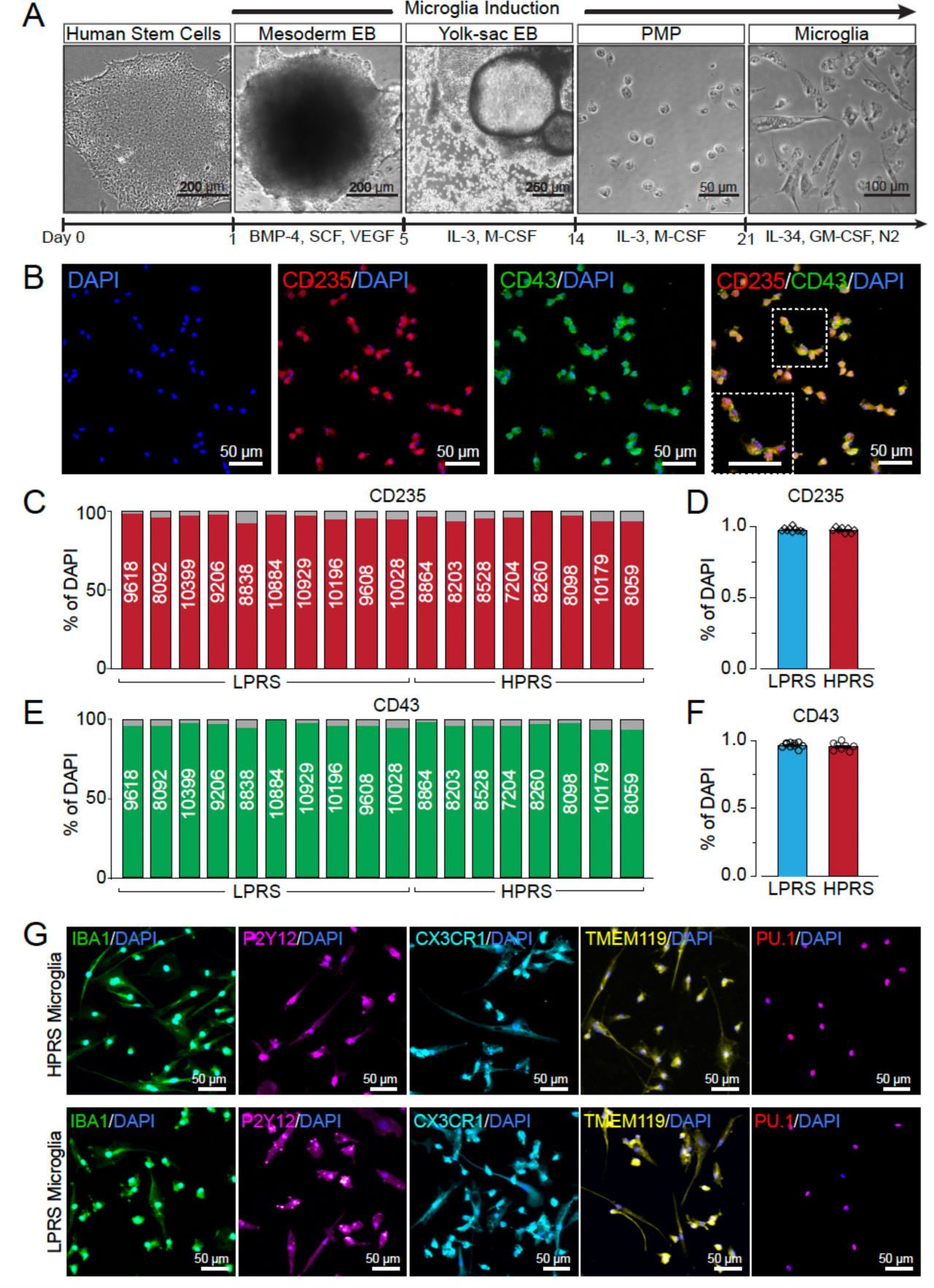
Characterization of iPSC-derived PMPs and microglia. **(A)** Protocols for generation of PMPs and microglial cells from iPSC. **(B)** Representative images of CD235^+^ and CD43^+^ cells in PMPs (line 8864). **(C, D)** Quantification of CD235^+^ PMPs derived from all iPSC lines with different PRS (LPRS, n=10; HPRS, n=8). **(E, F)** Quantification of CD43+ PMPs derived from all iPSC lines with different PRS (LPRS, n=10; HPRS, n=8). **(G)** Representative images of IBA1+, P2Y12+, CX3CR1+, TMEM119+, and PU.1+ microglia from HPRS line 8864 and LPRS line 9618.

### Differential transcriptomic profiles between HPRS and LPRS microglial cells

We conducted bulk RNAseq analysis in human microglia generated from iPSCs (*45*) (n=9, 4 HPRS, and 5 LPRS, both sexes, **Table s1**). We first compared the expression levels with a publicly available single-cell RNA-Seq data set of adult human brain cells (*46*). The iPSC-derived microglia and adult human microglial cells show very similar transcriptomic profiles (**Fig. s1C**), indicating the appropriate differentiation of our human microglia.

We identified 1,980 differentially expressed genes (DEGs) between HPRS and LPRS microglial cells in the absence of ethanol (**Fig. 2A**). Among these DEGs, the genes with higher expression (n = 1013) in AUD HPRS showed significant enrichment of GO biological process terms related to chromosome separation regulation and mitotic sister chromatid separation (**Fig. 2B**). Additionally, these genes were significantly enriched for receptor activator (**Fig. 2C**) and cellular component ontologies relating to chromosomes and centromeric regions (**Fig. 2D**). On the other hand, the genes with lower expression (n = 967) in AUD HPRS exhibited enrichment in biological functions related to immune signaling, specifically involving peptide antigen assembly with the MHCII protein complex and antigen processing and presentation (**Fig. 2B**). In terms of molecular function, these lower DEGs were predominantly linked to immune receptor activity, antigen binding, and MHCII protein complex binding (**Fig. 2C**), with their cellular component enrichment observed in the MHCII protein complex (**Fig. 2D**). Notably, a higher expression level of “chromosome separation” genes in the HPRS microglial cells was observed when compared with LPRS microglial cells (**Fig. 2E**). Also, we noticed lower expression levels of genes belonging to the “antigen processing and presentation” terms in AUD LPRS cells (**Fig. 2F**). There was, as expected, considerable variability among individual lines.

**Fig. 2.**
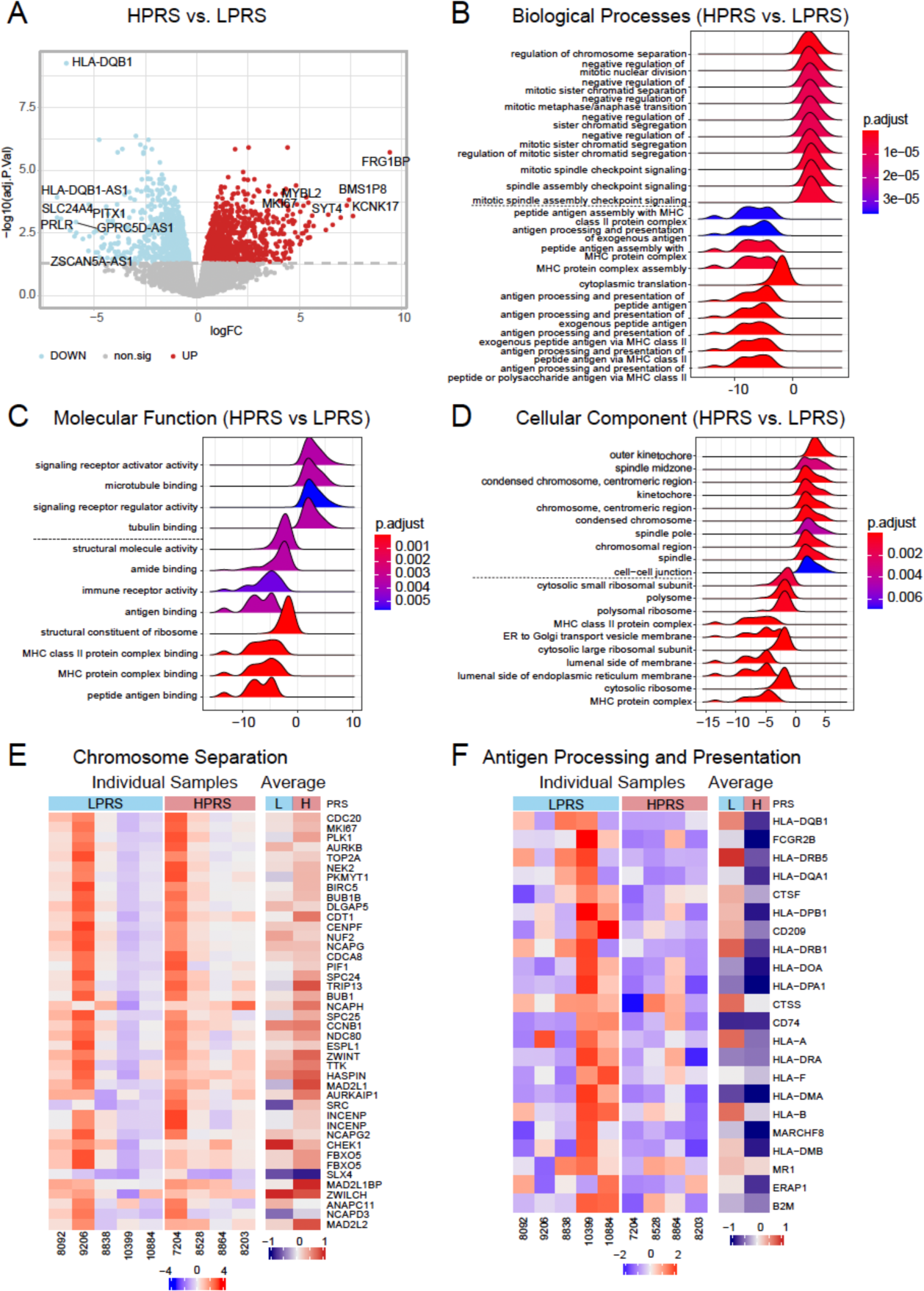
Whole genome transcriptome profile comparison between human microglial cells derived from HPRS and LPRS iPSC lines. **(A)** Volcano plot depicts the differential expressed genes (DEGs) between the HPRS and LPRS microglial cells. **(B-D)** Gene Ontology (GO) analysis depicting the significant terms (top 10 up- and down-regulated terms) associated with microglia when comparing HPRS vs. LPRS. It includes biological processes (BP, B), molecular functions (MF, **C**), and cellular components (CC, **D**). P.adjust values are displayed. **(E)** The heatmap illustrates gene expression patterns related to chromosome separation in both HPRS (red, n=4 lines) and LPRS (blue, n=5 lines) microglia. **(F)** The heatmap illustrates gene expression patterns related to antigen processing and presentation in both HPRS (red, n=4 lines) and LPRS (blue, n=5 lines) microglia.

### Activation of human microglial cells in response to ethanol exposure

To gain insight into the relationship between AUD PRS and ethanol exposure in microglia, we next investigated whether ethanol exposure causes differential cellular responses in human HPRS (n=5) vs LPRS (n=6) microglial cells (**Table S1**). We used intermittent ethanol exposure for 7 days, with ethanol replenished daily, as reported previously (*47*). It has been reported that ethanol activates microglial cells in rodents (*48, 49*). To investigate if ethanol activates human microglia in culture, we assessed the expression of CD68, a transmembrane protein found in plasma, lysosomal, and endosomal membranes that indicates microglial activation (*50*). In both HPRS and LPRS human microglial cells, we observed elevated expression of CD68 after ethanol exposure (**Fig. 3A&B**). The level of ethanol induced CD68 expression was similar between HPRS and LPRS microglial cells (**Fig. 3A&B**). Since activated microglia undergo morphological changes from a “ramified” state, a highly branched and elongated appearance, to an “amoeboid” state, with retracted processes and a more rounded shape (*51*), we used “Fractal analysis” in ImageJ at the single-cell level to evaluate human microglial morphology. (**Fig. 3C, Fig. S1G**). In the absence of ethanol, HPRS microglial cells displayed lower fractal dimension values and higher circularity compared to LPRS microglial cells (**Fig. 3D, Fig. S1H-J**). Following ethanol exposure, there was a reduction in fractal dimension observed in both HPRS and LPRS microglial cells (**Fig. 3D**), indicating a decrease in the complexity of microglial morphology in response to ethanol exposure. Notably, HPRS microglial cells exhibited a more significant decrease in fractal dimension and spine ratio, alongside a marked increase in circularity relative to their LPRS counterparts after exposure to ethanol. These observed differences were notably more substantial under the 75mM ethanol condition (**Fig. 3D**). These data indicated that human microglial cells were activated by ethanol exposure, and there are differences in morphological features between HPRS and LPRS microglial cells, suggesting genetic background may affect cellular responses to ethanol.

**Fig. 3.**
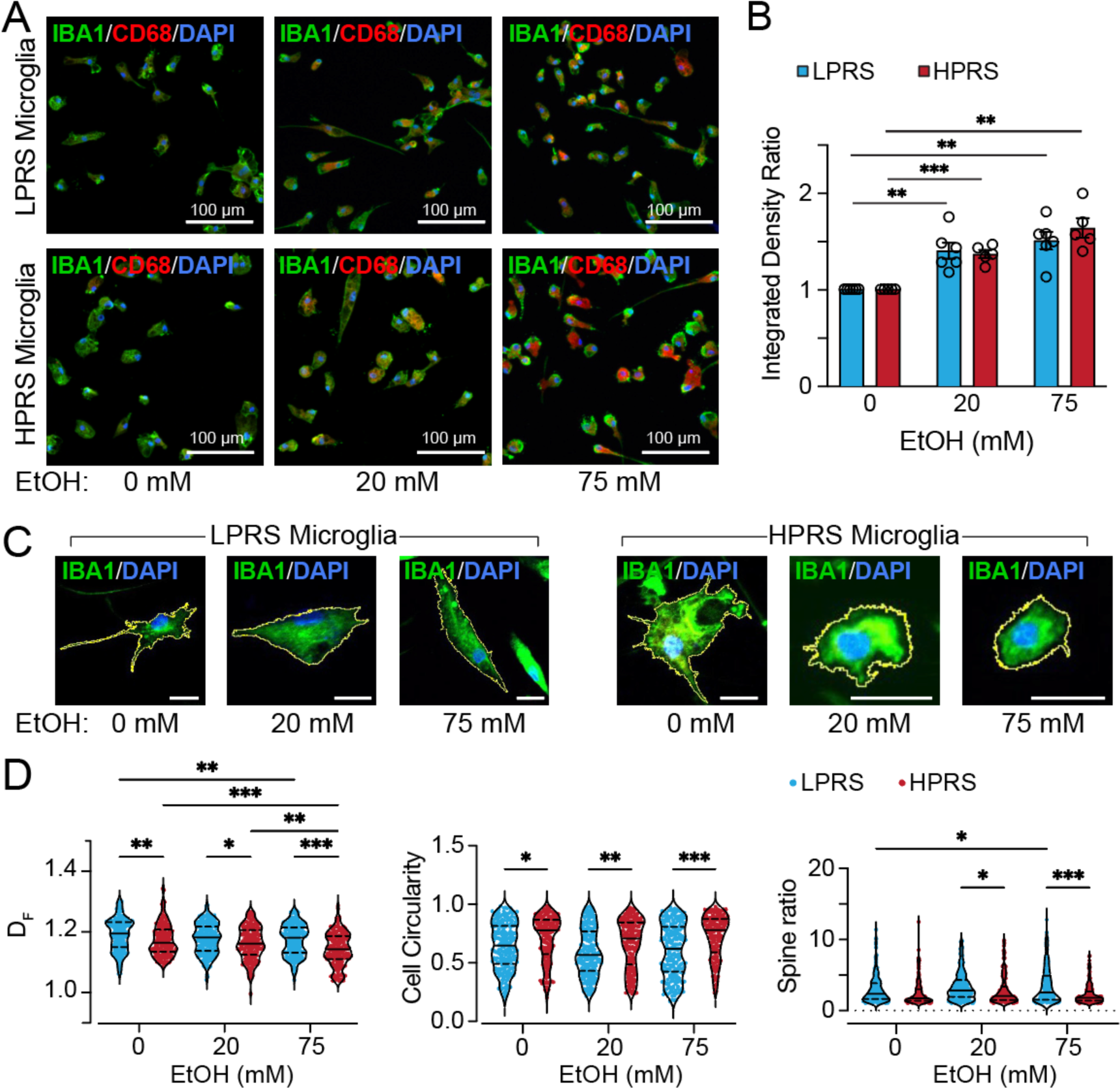
The impact of Ethanol exposure on the activation of iPSC-derived microglia. **(A)** Representative images of CD68^+^ and IBA1^+^ microglia derived from HPRS (line 8528) and LPRS lines (line 8206) following 7-day exposure to Ethanol (0 mM, 20 mM, and 75 mM). **(B)** Quantification of CD68 corrected fluorescence integrated density normalized to each control (0 mM) replicate. LPRS (n=6 lines) and vs. HPRS (n=5 lines), ** < 0.01, *** < 0.001. Data are presented as mean ± SEM. **(C)** Representative images showing the process traced with regions of interest (ROI) using the IBA1 signal in iPSC-derived microglia with HPRS (line 8864) or LPRS (line 8838). **(D)** Morphological analysis of microglia binary outlines using FracLac ImageJ. n = 177 cells/ 5 HPRS lines, 211 cells/ 6 LPRS lines for 0 mM; n = 188 cells/ 5 HPRS lines, 214 cells/ 6 LPRS lines for 20 mM; n = 164 cells/ 5 HPRS lines, 219 cells/ 6 LPRS lines for 75 mM. * < 0.05, ** < 0.01, *** < 0.001. Data are presented as medium ± quartiles.

### Transcriptomic profile differences of HPRS and LPRS microglial cells in response to ethanol exposure

We next determined the effects of intermittent ethanol exposure on the transcriptomic profiles of the iPSC-derived microglia (n=9, 4 HPRS, and 5 LPRS, **Table S1**). Exposure to ethanol did not affect human microglial cell identity, as determined by comparing our microglial bulk RNA-Seq data with the publicly available single-cell RNA-Seq data set of adult human brain cells (*46*) (**Fig. S1C**). Interestingly, some *ADH* and *ALDH* genes, including *ADH5*, *ALDH1A1*, *ALDH1A2*, and *ALDH2*, which encode enzymes involved in metabolizing ethanol into acetaldehyde and acetate (*52*), were found to be expressed by human microglia (**Table S2**), suggesting they may metabolize ethanol.

To investigate the different impacts of intermittent ethanol exposure on the transcriptomic profiles, we compared gene expression profiles with and without ethanol exposure (0mM vs, 75mM) (**Table S1**). We identified 3169 DEGs associated with intermittent ethanol exposure in HPRS microglial cells and 2472 DEGs in LPRS microglial cells (**Fig. 4A&B**). Among these DEGs, 1601 were upregulated in HPRS microglial cells, while 1308 were upregulated in LPRS microglial cells after intermittent ethanol exposure, with 909 genes overlapping between HPRS and LPRS microglial cells (**Fig. S2**). GO analysis unveiled significant enrichment of terms from the upregulated DEGs in HPRS lines, but not in LPRS lines, in biological processes related to peptide antigen assembly with the MHCII protein complex and antigen processing and presentation (**Fig. 4C**&**D**). Subsequently, we plotted running scores from GSEA explicitly focusing on the GO biological process terms unique to HPRS microglial cells. In addition to the terms related to antigen processing and presentation via the MHCII protein complex, this analysis uncovered significant enrichment of terms related to the regulation of phagocytosis (**Fig. 4E&F**). This observation underscores the impact of ethanol on antigen presentation, a critical process closely tied to phagocytosis, which encompasses the uptake, processing, and presentation of antigens to initiate adaptive immune responses.

**Fig. 4.**
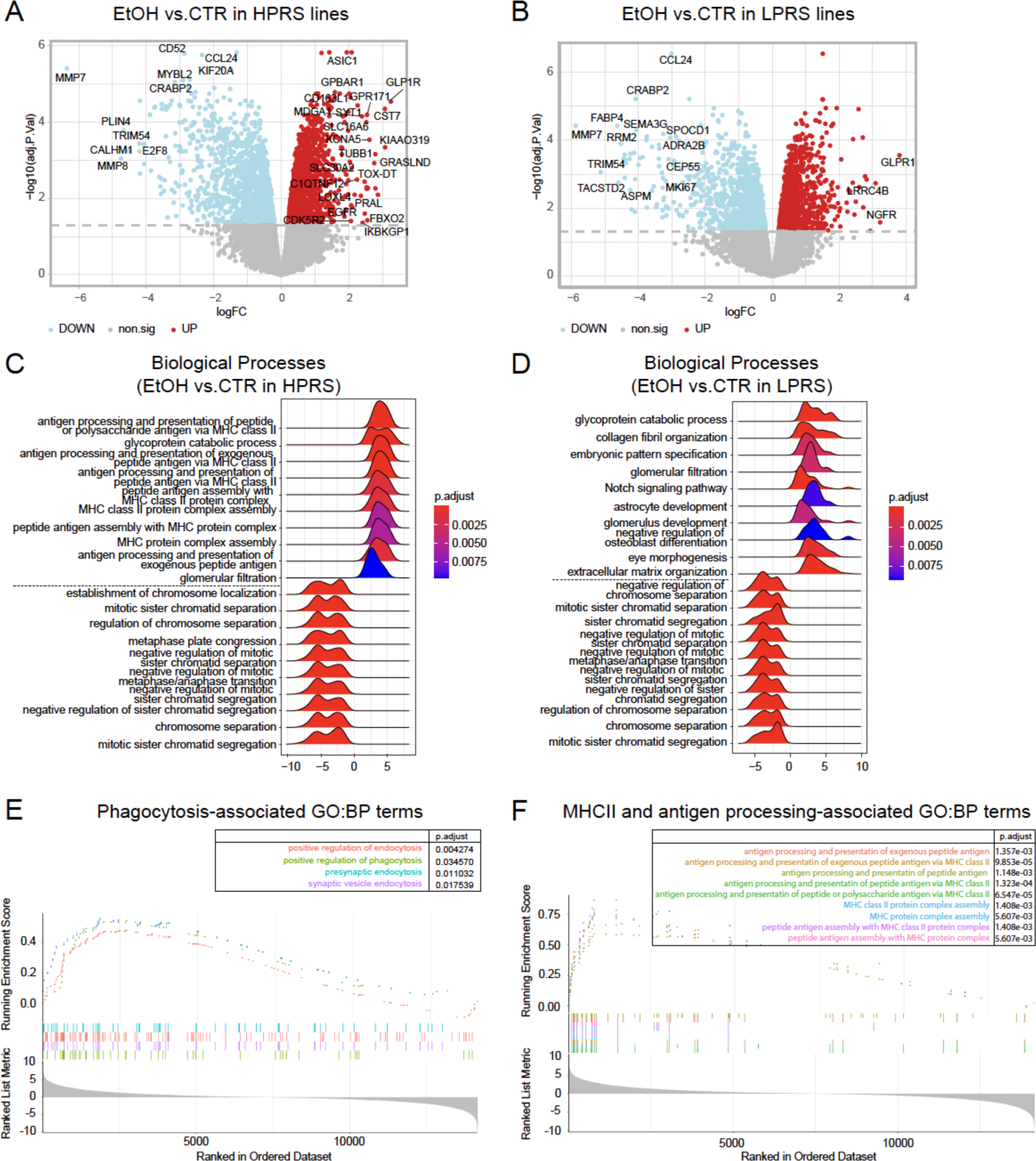
Transcriptomic profiles in HRPS vs. LPRS human microglia after ethanol exposure. **(A-B)** Volcano plots depict the differentially expressed genes (DEGs) between the different Ethanol concentrations (75mM vs. 0mM) in 4 HPRS lines (A) and 5 LPRS lines (B) of microglial cells. **(C-D)** Gene Ontology (GO) analysis of biological processes (BP) highlighting significant terms (top 10 up- and down-regulated terms) associated with ethanol treatment (75 mM vs. 0 mM) in both HPRS (C) and LPRS (D) microglia. Adjusted P-values (P.adjust) are displayed. **(E)** GSEA plot depicting the enrichment of the phagocytosis-associated pathway specific to HRS microglia before and after Ethanol treatment. P-values and P.adjust are displayed. **(F)** GSEA plot of the MHCII and antigen processing associated pathway enrichment specific to HRS microglia before and after Ethanol treatment. P-values and P.adjust are displayed.

Genes related to antigen processing and presentation are shown in **Fig. 5A**, and genes associated with phagocytosis in **Fig. 5C**. Most of these genes showed a significant increase in expression in HPRS microglial cells following exposure to ethanol. In contrast, their expression levels remained relatively stable in LPRS microglial cells (**Fig. 5B&D**). Notably, one of the genes identified in this category, *CLEC7A* (Dectin-1, encoding a mammalian C-type lectin receptor expressed on microglial cell surfaces that is known to promote zymosan phagocytosis (*53*)), exhibited substantial upregulation in HPRS microglial cells after ethanol exposure (**Fig. 5D**). To validate these findings, we performed immunohistochemistry to assess the expression of CLEC7A protein by immunostaining in each of the 18 cell lines individually (**Fig. 6A, Table S1**). In agreement with mRNA measurements, we found a notable increase in CLEC7A immunoreactivity expression levels in HPRS microglial cells following ethanol exposure (75mM). In contrast, the levels remained relatively stable in LPRS microglial cells (**Fig. 6B, C**).

**Fig. 5.**
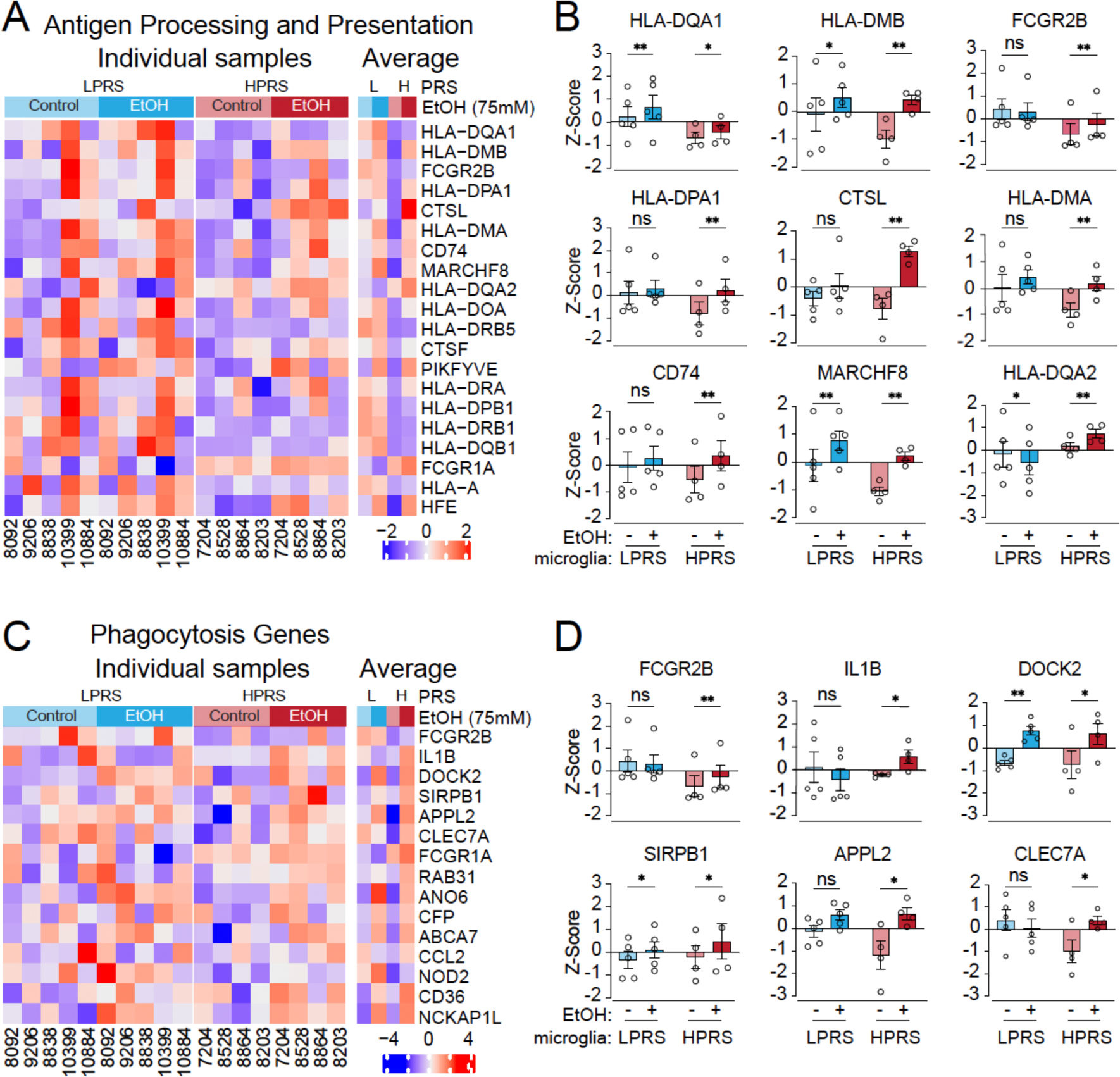
Differential expression of phagocytosis-related genes between HPRS and LPRS microglia in response to ethanol exposure. **(A)** A heatmap illustration of the gene expression profiles related to antigen processing and presentation in both HPRS (pink) and LPRS (blue) microglia, both before (green) and after (red) Ethanol treatment. **(B)** Box blots comparisons of the differential expression of genes associated with antigen processing and presentation in microglia from both HPRS (pink) and LPRS (blue) lines before and after Ethanol treatment. LPRS/HPRS lines = 5/4. * < 0.05, ** < 0.01. Data are presented as mean ± SEM. **(C)** A heatmap illustration of the gene expression profiles related to the phagocytosis pathway in both HPRS (pink) and LPRS (blue) microglia, both before (green) and after (red) Ethanol treatment. **(D)** Box blots comparison of the representative differential expression of genes associated with the phagocytosis pathway in microglia from both HPRS (pink) and LPRS (blue) lines before and after Ethanol treatment. LPRS/HPRS lines = 5/4. * < 0.05, ** < 0.01. Data are presented as mean ± SEM.

**Fig. 6.**
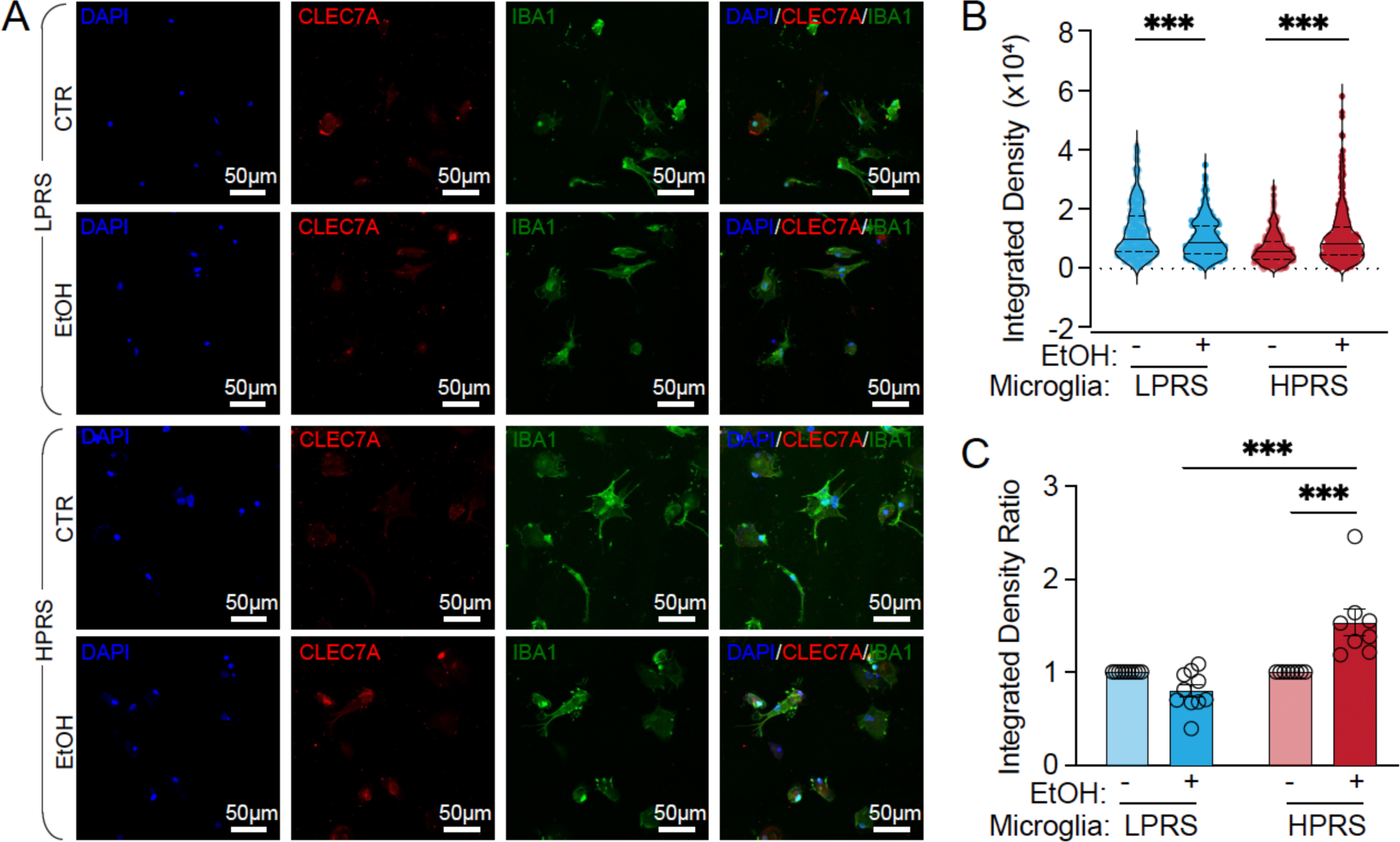
CLEC7A expression between HPRS and LPRS microglia after ethanol exposure. **(A)** Representative images of CLEC7A^+^ and IBA1^+^ microglia derived from PRS lines following ethanol exposure (0 mM and 75 mM). **(B, C)** Quantification of CLEC7A^+^ fluorescence integrated density **(B)** and corrected fluorescence integrated density normalized to each control (0 mM) replicate **(C).** n = 331 cells/8 HPRS lines, and 365 cells/10 LPRS lines for 0 mM; n = 315 cells/8 HPRS lines, and 373 cells/10 LPRS lines for 75 mM; ** < 0.01, *** < 0.001. Data are presented as mean ± SEM.

Downregulated genes affected by intermittent ethanol exposure in HPRS and LPRS microglial cells have similar predicted functions. They are significantly enriched in cell cycle regulation, including chromosome separation and mitotic sister chromatid separation **(Fig. 4C&D)**. To better visualize the gene expression changes in ethanol-responsive DEGs and cell cycle phases in both LPRS and HPRS microglial cells, we generated a heatmap summarizing the expression of cell cycle-related genes **(Fig. S3A)**. Upon ethanol exposure, the pathway analyses for the upregulated genes showed enrichment in the G1/S and G2/M DNA damage checkpoint markers **(Fig. S3B, Fig. S4)**. This suggests that ethanol may cause cell cycle arrest and/or DNA damage in HPRS and LPRS microglial cells. An increase in γ-H2AX (a sensitive molecular marker of DNA damage (*54*)) after exposure to 75 mM ethanol was found by immunostaining in both HPRS and LPRS microglial cells **(Fig. S3C&D)**. Consistent with a previous report in mouse neurons (*55*), these results indicated that intermittent ethanol exposure may cause DNA damage in human microglia, potentially interfering with microglial proliferation.

### Differential phagocytic function in HPRS and LPRS human microglial cells after intermittent ethanol exposure

As noted above, HPRS and LPRS microglial cells showed differential morphological changes and differential gene expression, particularly upregulation of genes related to phagocytosis, in HPRS microglial cells following intermittent ethanol exposure. Thus, we hypothesized that HPRS and LPRS would show differential phagocytosis after intermittent ethanol exposure. To test this hypothesis, we utilized pH-sensitive pHrodo zymosan beads to assess the phagocytic activity (*56, 57*) in microglial cells derived from 18 lines individually, both with HPRS and LPRS, after 7-day intermittent ethanol exposure at two different ethanol concentrations (20 mM, and 75 mM; **Fig. 7A&B**). At baseline (ethanol-free), we did not observe a significant difference between HPRS and LPRS in the integrated density of zymosan beads within the microglial cells (**Fig. 7C**). However, we did notice a higher proportion of microglia containing beads in LPRS microglial cells compared to HPRS microglial cells without ethanol exposure (**Fig. 7D**). Following intermittent ethanol exposure, we observed significantly augmented phagocytosis in HPRS microglial cells, particularly at higher ethanol concentrations (75mM) (**Fig. 7B&C**), while LPRS cells did not exhibit a corresponding increase (**Fig. 7A-C**). A similar pattern was observed in the proportion of microglia containing beads, which substantially increased in HPRS microglial cells but remained relatively unchanged in LPRS microglial cells after intermittent ethanol exposure (**Fig. 7D**). Our results indicated that ethanol exposure enhances phagocytic activity in HPRS, but not in LPRS microglial cells. These results suggested a potential link between variation in genetic background summarized by PRS and ethanol-induced alterations in microglial function.

**Fig. 7.**
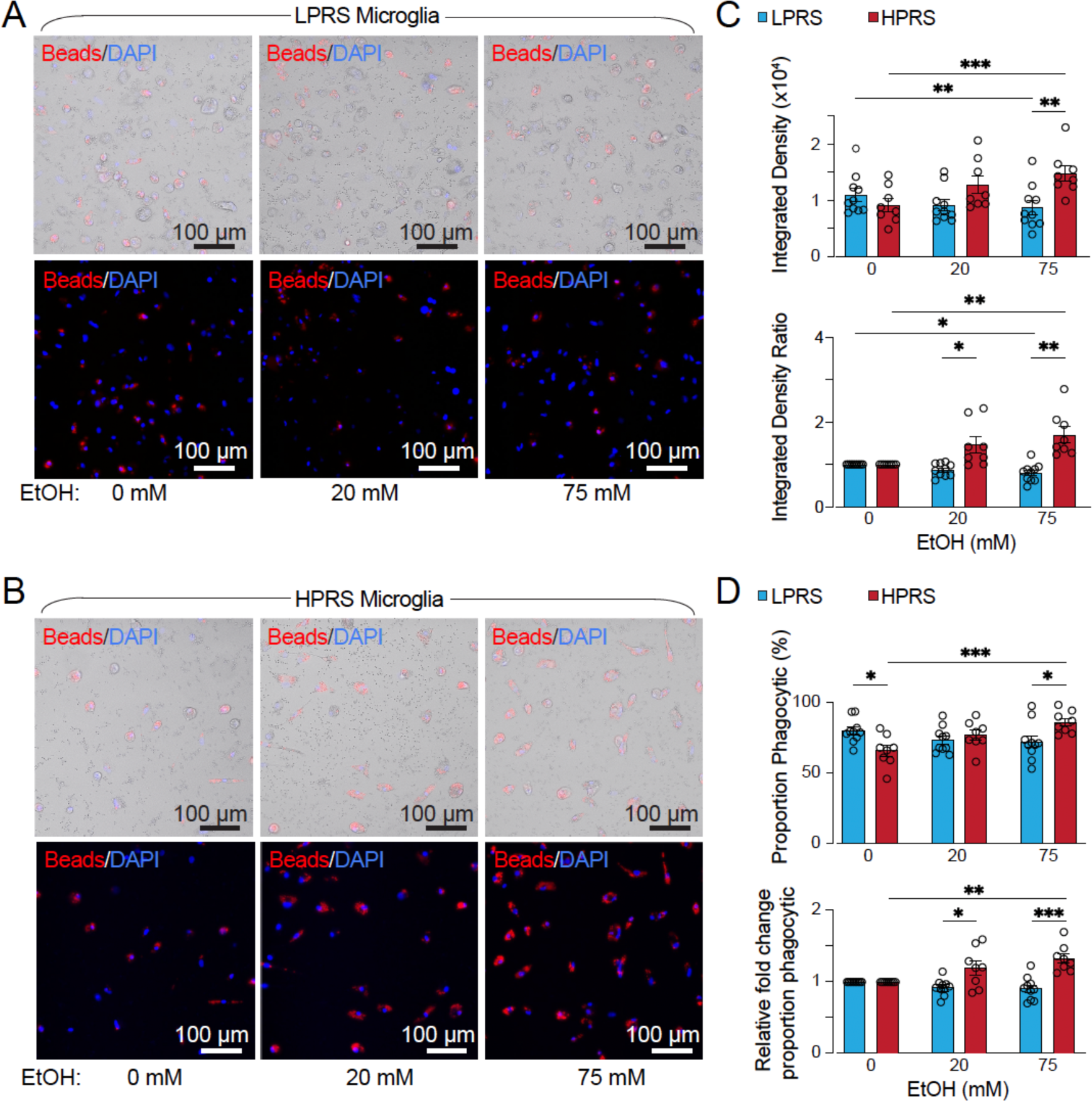
Enhanced phagocytosis ability in HPRS iPSC-derived microglia in response to Ethanol. **(A, B)** Representative live-cell imaging of iPSC-derived microglia from LPRS (line 8092, A) and HPRS (line 8528, B) engaged in the phagocytosis of zymosan particles at different ethanol concentrations (0 mM, 20 mM, and 75 mM). The upper images are overlapped with bright-field microscopy for reference. **(C)** Quantification of the fluorescence-integrated density of zymosan bioparticles (beads): the upper panel shows the fluorescence-integrated density between HPRS and LPRS microglial cells, while the lower panel shows the corrected fluorescence-integrated density normalized to each control (0 mM). LPRS/HPRS lines = 10/8, ** < 0.01, *** < 0.001. Data are presented as mean ± SEM. **(D)** Quantification of the proportion of microglia with zymosan particles (beads): the upper panel shows the proportion of microglia with beads, while the lower panel displays the corrected proportion, which has been normalized to each control (0 mM) replicate. LPRS/HPRS lines = 10/8, ** < 0.01, *** < 0.001. Data are presented as mean ± SEM.

Microglia interact with neurons through synaptic pruning to shape synapse formation and synaptic transmission (*20, 21*). We thus examined the impact of AUD HPRS and LPRS microglial cells on human iPSC-derived neurons. We individually co-cultured each line of microglial cells (6 lines of AUD HPRS and 7 lines of AUD LPRS, see **Table S1** for detail) with induced human induced neuronal (iN) cells, using ectopic expression of the transcription factor neurogenin 2 (Ngn2) (*58*). The iNs were induced from an iPSC line with no diagnosed AUD, as used in a previous study (*59*). Thereby, we were able to evaluate the differential effects between HPRS and LPRS microglia on a common set of “neutral” neurons. To facilitate synapse formation, these iNs were cultured on monolayer mouse glial cells (*60, 61*).HPRS or LPRS microglial cells were added to iN cultures on day 14 post-induction. The final microglial density in culture (day 35-40 post-induction) is around 7-9% of total cell numbers, including mouse glial cells **(Fig. 8A&B)**, which is consistent with the microglia proportion in the human brain (*62*). We first analyzed synapse formation after 7-day intermittent ethanol exposure with 75 mM ethanol. After immunohistochemical staining with the presynaptic marker synapsin and the neuronal marker MAP2, we used *Intellicount* (*63*) to quantify the synaptic density per cell. As a control, we also included iN cultures (on mouse glia) without microglia. We found that synaptic density was increased in human iN cultures after intermittent ethanol exposure **(Fig. 8C&D).** While adding LPRS microglial cells had no substantial impact on the synaptic density, HPRS microglial cells reversed the increase of synapse number after intermittent ethanol exposure (**Fig. 8C&D**). To further validate the impact of HPRS microglial cells on synaptic formation/function, we conducted patch-clamp electrophysiology on iNs with or without microglial cells. Consistent with the findings of the synaptic density analyses, after intermittent ethanol exposure, we found increased frequencies of miniature excitatory postsynaptic currents (mEPSCs) in iNs without human microglia or with LPRS microglial cells but not in HPRS-iN co-cultures (**Fig. 8E&F**).

**Fig. 8.**
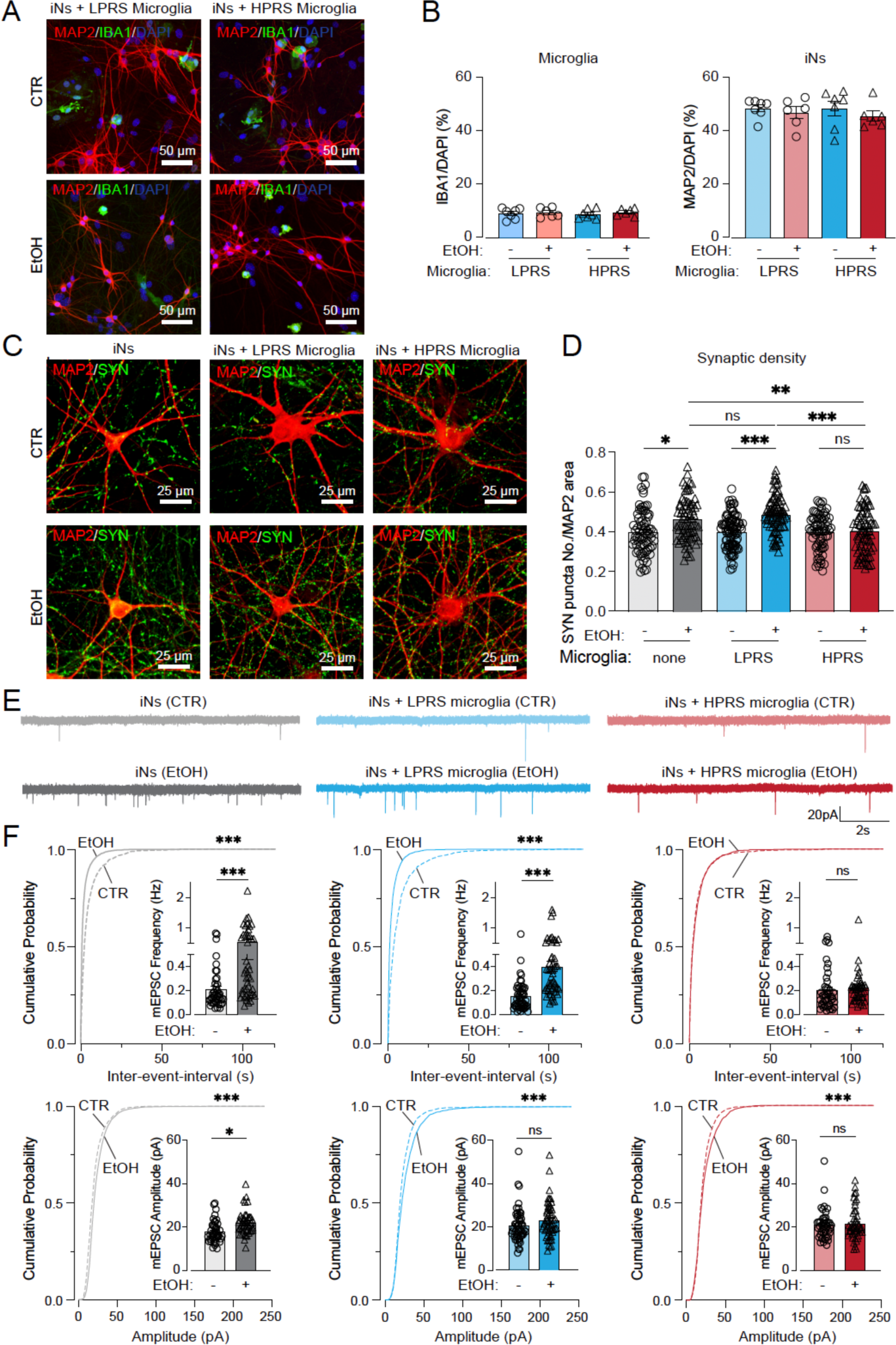
HPRS iPSC-derived microglia decreases excitatory neurotransmission in coculture. **(A)** Representative images of coculture consisting of MAP2^+^ neurons and IBA1^+^ microglia derived from both HPRS and LPRS lines, following treatment with ethanol (0 mM and 75 mM). **(B)** Quantification of the proportion of microglia and neurons in coculture with HPRS and LPRS microglia-ngn2, under 0mM and 75mM conditions (LPRS = 7 lines, HPRS = 6 lines). Data are presented as mean ± SEM. **(C)** representative images illustrating synaptic puncta (green, labeled by synapsin immunofluorescence) associated with dendrites (red, visualized by MAP2 immunofluorescence) in iNs without microglia, with LPRS microglial cells, or with HPRS microglial cells. **(D)** Quantification of synapsin puncta densities per MAP2 area. (iN without microglia: 0mM = 73 images, 75mM = 73 images; iN + LPRS microglial cells: 0mM = 93 images, 75mM = 85 images; iN + HPRS microglial cells: 0mM = 70 images, 75mM = 75 images). **(E)** Representative traces of miniature excitatory postsynaptic currents (mEPSCs) in Ngn2-iNs without microglia (grey, 0mM = 43 cells, 75mM = 46 cells), Ngn2-iNs with LPRS microglial cells (blue, 0mM = 53 cells, 75mM = 55 cells), and Ngn2-iNs with HPRS microglial cells (red, 0mM = 43 cells, 75mM = 46 cells). **(F)** Cumulative distribution and quantification (inserts) of mEPSC frequency (upper panel) and amplitude (lower panel) in Ngn2-iNs without microglia (grey N), Ngn2-iNs with LPRS microglial cells (blue), and Ngn2-iNs with HPRS microglial cells (red). Data are mean ± SEM; Statistical significance (*p < 0.05, **p < 0.01, ***p < 0.001) was evaluated with the Kolmogorov–Smirnov test (cumulative probability plots) and t-test (bar graphs).

Together, these findings suggest that ethanol partially induced adaptive changes in the strength of excitatory transmission by increasing synapse numbers in human neurons. AUD HPRS microglial cells may attenuate the impact of these phenotypic changes through heightened synaptic pruning/elimination activity after ethanol exposure.

## Discussion

AUD is a complex genetic disorder, and although most individual genetic variants do not have genome-wide significant effects, when considered collectively in a PRS, they quantify part of an individual’s susceptibility to developing the disorder (*64-66*). We investigated how the polygenic background, represented as PRS, affects microglial function, how intermittent exposure to ethanol affects microglial function, and how the two factors interact. It is likely that many of the genetic variants summarized in PRS can be predicted to affect gene regulatory networks either directly or indirectly. In the absence of ethanol, we found significant differentially expressed genes when comparing microglia from individuals with AUD and a high PRS for AUD (HPRS) to those from individuals without AUD and with low PRS (LPRS). The genes more highly expressed in HPRS microglial cells were enriched in pathways related to receptor activation and chromosome separation, while those expressed at lower levels were enriched in genes related to immune signaling, particularly with the MHCII complex.

Following Intermittent ethanol exposure, we found that HPRS and LPRS microglial cells rapidly transition from their default branched profile (*67*) to a more amoeboid state (*68*). Microglia with HPRS showed a significant decrease in fractal dimension and a pronounced increase in circularity compared to their LPRS counterparts after ethanol exposure. Further gene expression profiling revealed ethanol exposure alters significant DEGs in HPRS and LPRS microglial cells, with more than half of the DEGs overlapping. The upregulated genes in HPRS were enriched for processes related to peptide antigen assembly with the MHCII protein complex and antigen processing and presentation, and phagocytosis. Microglia, functioning as antigen-presenting cells (APCs), are equipped with phagocytic receptors that facilitate the capture of antigens. These antigens undergo processing within phagosomes and are subsequently presented by MHC class II and costimulatory molecules expressed in microglia (*69, 70*). The whole process could modulate the immune responses within the central nervous system (CNS) during AUD.

Among those upregulated genes, we found significant changes in CLEC7A in HPRS microglial cells at both the transcript and protein levels following exposure to ethanol. CLEC7A is a receptor located on the surface of microglial cells which plays a crucial role in the immune response by detecting zymosan, a type of fungal β-glucan, and subsequently initiating the phagocytic process in microglia (*53, 71, 72*). This heightened CLEC7A expression in HPRS microglial cells may partially explain the increased phagocytic activity following exposure to ethanol. Additionally, studies have identified CLEC7A as a prominent member of the disease-associated microglia (DAM) genes (*73*), consistently showing elevated expression by microglia in diverse mouse models of neurodegeneration, including Alzheimer’s disease (AD) models (*74, 75*). Thus, our observation may also suggest a potential connection between the elevated CLEC7A in HPRS microglial cells and AUD-related dementia. In a recent study, they observed notable differences in the expression of genes associated with human AD among AD mice with various genetic backgrounds. Specifically, the B6.APP/PS1 and WSB.APP/PS1 strains displayed higher expression levels for the gene Clec7a, while the CAST and PWK strains did not (*76*). These results suggest that genetic background may modulate microglial phagocytic function in response to ethanol and influence neuron function in AUD, potentially affecting the interaction between microglia and neuronal cells in the context of AUD pathophysiology.

Microglia can modulate synapses through processes such as partial phagocytosis of synaptic segments or engulfing entire pre- or postsynaptic elements (*77, 78*). After observing that ethanol can promote microglial phagocytosis of zymosan in HPRS lines, we began to investigate the potential effects of HPRS microglial cells on synaptic transmission in the context of alcohol consumption. To explore this further, we established a coculture system incorporating human Ngn2-iNs and microglia. Our findings indicated that ethanol enhances excitatory synaptic transmission, which aligns with previous research demonstrating increased glutamatergic activity following chronic alcohol exposure (*79*). While we did not observe significant changes in synaptic densities in neurons co-cultured with microglia compared to neurons without microglia, we did notice that HPRS microglial cells may contribute to a reduction in excitatory synaptic densities and transmission in neurons following intermittent ethanol exposure. This suggests that HPRS microglial cells may play a crucial role in shaping synaptic connectivity through a process known as microglia-dependent synapse elimination specifically in HPRS individuals, potentially rendering them at higher risk for developing/maintaining AUD. A mouse study (*28*) has shown that long-term alcohol exposure activates microglia to phagocytose synapses, leading to impaired synaptic plasticity and cognitive function (*48*). Consequently, our results imply that the heightened phagocytic activity observed in HPRS microglial cells after ethanol exposure may exert a significant influence on synaptic connectivity and remodeling, potentially impacting neuronal function in the context of alcohol use disorder-related dementia.

It remains to be explored how ethanol exposure regulates gene expression in human microglial cultures and how AUD PRS influences the gene expression and functionality of microglial cells. Alcohol can also directly activate microglial cells to increase the production of proinflammatory chemokines and cytokines, which may regulate gene expression in microglial cells (*80*). Neurons can metabolize ethanol into acetate (*55*), but whether microglial cells can produce acetate and highly reactive acetaldehyde from ethanol remains to be investigated. Nevertheless, human microglial cells produced from iPSCs express multiple ADH and ALDH genes necessary to participate in the metabolic process for ethanol (**Table S2**). The ethanol metabolite acetate can be a significant source of acetyl-CoA produced by chromatin-bound acetyl-CoA synthetase, essential for histone acetylation and epigenetic regulation for gene expression (*81*). It is also possible that the excessive alcohol consumption in the AUD HPRS subjects that the HPRS iPSCs were derived from carried certain epigenetic changes that confer the differences between gene expression and functional differences between the HPRS vs. LPRS microglial cells. How AUD and PRS may interact with histone acetylation, as affected by ethanol metabolites and open chromatin state to influence gene expression in vivo and in vitro, needs to be further elucidated.

It is important to acknowledge certain limitations in our study. Notably, our experimental approach relied on a two-dimensional cell culture, which, while valuable for investigating microglial responses and gene expression patterns, does not recapitulate the complex three-dimensional interactions that occur in the brain. Our analysis of microglial-neuron interactions utilized only excitatory neurons, limiting the interpretation of our findings. Further investigations employing brain organoid models (*45*) or in vivo (*30*) studies will help gain a more comprehensive understanding of the dynamics and functional consequences of microglia-neuron interactions in AUD and the role of genetic factors in these interactions.

In summary, we observed differential gene expression patterns in HPRS and LPRS human microglial cells in response to intermittent ethanol exposure and heightened phagocytotic responses in the AUD HPRS microglial cells that may impact synaptic transmission differentially than LPRS microglial cells. These findings shed light on the multifaceted mechanisms underlying AUD and emphasize the importance of considering genetic factors when exploring microglial responses in this complex disorder.

## Materials and Methods

### Human induced pluripotent stem cell (iPSC) lines

The human iPSC lines utilized in this study were derived from lymphocytes and lymphoblastoid cells collected from the Collaborative Study on the Genetics of Alcoholism (COGA) study participants obtained from the NIAAA/COGA Sharing Repository. The selection criteria for these individuals were based on previously established PRS for AUD (*12*). Specifically, to investigate PRS for AUD, we selected 8 individuals diagnosed with AUD according to the Diagnostic and Statistical Manual, fifth edition (*82*) with PRS greater than the 75th percentile and 10 individuals who had no history of AUD and exhibited PRS lower than the 25th percentile (**Table S1**). Cryopreserved lymphocytes were thawed, and the erythroblastic cells expanded and infected with the Sendai virus, expressing reprogramming factors (CytoTune, Life Technologies). All reprogramming and assessment of pluripotency was performed by Sampled, formerly Infinity BiologiX LLC. Cultures of iPSC were maintained feeder-free on Matrigel in mTeSR+ medium (STEMCELL Technologies) and routinely passaged with Accutase (STEMCELL Technologies).

### Differentiation and culture of human iPSC-derived primitive macrophage progenitors (PMPs) and microglia

PMPs were derived from iPSC lines using an established protocol (*45*). To initiate the formation of yolk sac embryoid bodies (YS-EBs), we exposed them to mTeSR+ medium (STEMCELL Technologies, 85850) supplemented with bone morphogenetic protein 4 (BMP4, 50 ng/ml), vascular endothelial growth factor (VEGF, 50 ng/ml), and stem cell factor (SCF, 20 ng/ml) for six days. For the induction of myeloid differentiation, YS-EBs were plated onto 10cm dishes containing X-VIVO 15 medium (Lonza) supplemented with interleukin-3 (IL-3, 25 ng/ml) and macrophage colony-stimulating factor (M-CSF, 100 ng/ml) (*34, 83*). Human PMPs emerged in the supernatant after four weeks post-plating and maintained production for over six months.

To achieve full maturation of microglia, we subjected the PMPs to a carefully designed 7-day treatment regimen. This involved the addition of interleukin-34 (IL-34, 100 ng/ml) and granulocyte-macrophage colony-stimulating factor (GM-CSF, 10 ng/ml) to DMEM/F12 (HyClone™, SH30023.01) containing N-2 supplement (Gibco™, 17502048). This protocol facilitated the maturation process of the PMPs into microglia.

### Generation of co-culture of iPSC-induced neurons (iNs) and iPSC-derived microglia

We utilized a mid-PRS line (line 420 (*59*)) iPSC sourced from the NIAAA/COGA Sharing Repository to generate excitatory induced neurons (Ngn2-iNs), following the protocol as described by (*58*).In brief, when the iPSCs reached a confluency of approximately 70%, they were dissociated using Accutase (Stem Cell Technologies) and subsequently plated onto a six-well plate coated with Matrigel® Matrix (Corning Life Sciences). These plated cells were then cultured in mTeSR plus™ media, which contained RHO/ROCK pathway inhibitor Y-27632, doxycycline-inducible lentiviruses expressing transcription factors (Ngn2), and the reverse tetracycline-controlled transactivator (rtTA). The cells were cultured at 37°C with 5% CO2. After around 6-8 hours of lentivirus infection, the culture medium was replaced with Neurobasal™ medium (GIBCO by Life Technologies). This medium was supplemented with B27, L-glutamine, and 2 mg/mL of doxycycline (MP Biomedicals) to induce TetO expression.

The lentivirus generation protocol was previously outlined by Wang et al. (*84*). Selection with puromycin (1 mg/mL; Sigma Aldrich, catalog# P8833) was carried out for the next 2 days. On day 5, the Ngn2-iNs (approximately 200,000 iNs per coverslip) were dissociated using Accutase™ and plated on glass coverslips coated with a monolayer of passage three primary mouse astrocytes (approximately 50,000 cells per coverslip) obtained from postnatal day 1-3 pups, following the procedure as described (*85, 86*). On day 6, fresh Neurobasal™ medium containing B27, L-glutamine, Ara-C (4 mM; Sigma Aldrich, catalog# C6645), 10 ng/mL of BDNF, NT3, and GDNF (Peprotech™) were added. Subsequently, every 5 days, starting from day 6, 50% of the culture medium was replaced with fresh Neurobasal™ medium supplemented with B27, L-glutamine, BDNF, NT3, and GDNF.

On day 14, iPSC-derived microglia (approximately 30,000 cells per coverslip, **Table s1**) were added into the Ngn2-iNs using fresh Neurobasal™ medium containing B27, L-glutamine, BDNF, NT3, GDNF, IL-34, and GM-CSF. Molecular, morphological, and functional analyses were conducted approximately 5 weeks after the initial induction with doxycycline.

### Intermittent ethanol exposure

Following a seven-day maturation period in DMEF/12 medium supplemented with N2, interleukin-34 (IL-34), and granulocyte-macrophage colony-stimulating factor (GM-CSF), human-derived microglia were subjected to a seven-day chronic intermittent exposure (CIE) paradigm using 200 Proof ethanol from Decon Laboratories (catalog# 2716) at concentrations of 0 mM (control), 20 mM, and 75 mM, in accordance with established protocols (*87-89*). Since ethanol is depleted in culture by evaporation (*47*), we replenished half the medium daily with the 20 mM or 75 mM ethanol throughout the intermittent ethanol exposure period. This meticulous daily renewal of the culture medium ensured continuous intermittent ethanol exposure, enabling the investigation of the effects of ethanol on these microglia.

### Immunohistochemistry (IHC)

For immunocytochemistry analysis, the cultured cells were fixed using freshly prepared buffered 4% paraformaldehyde. Subsequently, after blocking with 20% normal goat serum and permeabilization for 10 minutes with 0.2% Triton X-100 in PBS, the cell cultures underwent an antibody labeling procedure. Specifically, the cultures were incubated overnight at 4°C with the following primary antibodies which were diluted in blocking buffer: rabbit anti-CD235 (Invitrogen, 90757), mouse anti-CD43 (Invitrogen, 763493), chicken anti-human IBA1 (Aves Labs, IBA1-0020), mouse anti-human TMEM119 (Cell Signaling Technology, 41134S), rabbit anti-human P2RY12 (Sigma, HPA014518), rabbit anti-human PU.1 (Cell Signaling Technology, 2266S), rabbit anti-human CX3CR1 (Biorad, AHP1589), mouse anti-human CD68 (Invitrogen, 10987212), mouse anti-human CLEC7A (R&D Systems, MAB1859), and mouse anti-γH2AX (phospho-histone Ser139, Millipore Sigma, 05-636), chicken anti-human MAP2 (EMD Millipore, AB5543), mouse anti-human synapsin (SYSY,106011). After incubation with primary antibodies and subsequent PBS washes, the cultures were exposed to secondary antibodies labeled with Alexa Fluor 488, 546, or 633, as appropriate (anti-mouse and/or anti-rabbit and/or anti-chicken Molecular Probes), for 60 minutes at room temperature. To visualize cell nuclei, DAPI (0.3 μg/ml, Roche) was used for nuclear staining. After staining, the samples were rinsed with PBS and mounted for analysis. Microscopic images were captured using a confocal microscope (Zeiss LSM700) for further examination and data acquisition.

### Image analysis

All images were acquired using Z stack maximal projection in Zeiss Zen blue software. To analyze microglial morphology at the single-cell level, we used the “FracLac plugin” in ImageJ (https://imagej.net/ij/plugins/fraclac/FLHelp/Introduction.htm) after we collected the binary images of each microglial cell (*90*). Here, we use the parameters including fractal dimension (D), lacunarity, convex hull area (CHA), convex hull perimeter (CHP), cell circularity (CC), and convex hull span ratio to assess the complexity and circularity of microglial cells.

Fractal dimension (D) is the method for discerning various microglial shapes, ranging from simple round forms to intricate branched structures (*91, 92*). In this study, box-counting software was employed to enumerate the number of boxes encompassing any foreground pixels within the outlined images, which were progressively processed on grids of diminishing calibers (Figure s1D). Lacunarity indicates variations in the soma and additional morphological features. This parameter quantifies shape heterogeneity, both in terms of translation and rotation invariance (*92*). Lacunarity is computed using the box-counting software FracLac, representing the pixel mass distribution in microglia images. Convex hull perimeter (CHP) is the single outline of the convex hull expressed in microns. Convex hull area (CHA) refers to the measurement of the area enclosed by the smallest convex polygon, which is defined as a polygon with all interior angles smaller than 180 degrees, encompassing the entire cell shape. Cell circularity (CC) was calculated as (4π × cell area) / (cell perimeter)^2^. The circularity value of a circle is 1. The convex hull span ratio is determined by calculating the ratio between the major and minor axes of the convex hull.

The analysis of synaptic puncta density was performed on iNs at day 35. This analysis was conducted using the dendrite marker MAP2 and synaptic markers Synapsin-1. The quantification of correlated synaptic puncta size and intensity was performed using Intellicount (*63*).

### RNA Library Construction, Quality Control and Sequencing

Total RNA from iPSC-derived microglia (**Supplementary Table 2**) was extracted using RNeasy Plus kit (Qiagen). Library construction and sequencing were performed by Novogene. Messenger RNA was purified from total RNA using poly-T oligo-attached magnetic beads. After fragmentation, the first strand cDNA was synthesized using random hexamer primers, followed by the second strand cDNA synthesis using either dUTP for a directional library or dTTP for the non-directional library. The non-directional library was ready after end repair, A-tailing, adapter ligation, size selection, amplification, and purification. The library was prepared through end repair, A-tailing, adapter ligation, size selection, USER enzyme digestion, amplification, and purification steps. The library was checked with Qubit and real-time PCR for quantification and bioanalyzer for size distribution detection. Quantified libraries were pooled and sequenced on Illumina NextSeq500 platforms by Novogene. Approximately 40-60 million paired reads were generated for each sample (**Table S3**). Reads were aligned to the hg38 human reference genome by HISAT2 (v.2.1.0) (*93*). Transcript counts were extracted using the featureCounts function of the Rsubread package. Details of the bioinformatics analysis methods for RNA-Seq data are provided in the **Supporting Information.**

### Phagocytosis Assay

In the phagocytosis assay, human-induced pluripotent stem cell (iPSC)-derived microglia were seeded in a 96-well plate following a 7-day exposure to a intermittent ethanol exposure paradigm at varying concentrations (0 mM as the control, 20 mM, and 75 mM). To visualize the cell nuclei, Hoechst stain was diluted at a ratio of 1:1000 and added to the microglia, followed by incubation at 37°C for 10 minutes. Subsequently, zymosan bioparticles (Invitrogen™, P35364) were prepared by vortexing and then subjected to ultrasonication for 5 minutes. These zymosan bioparticles were then added to the live microglia in live cell imaging solution (A14291DJ), and the cells were incubated for 2 hours at 37°C. Live cell images were acquired using a BZ-x800 microscope equipped with Keyence BZ-x800 software for subsequent analysis and data collection.

### Electrophysiology

Functional analyses of iN cells were conducted using whole-cell patch-clamp electrophysiology as described elsewhere (*84, 94*). Miniature postsynaptic currents were recorded at a holding potential of −70 mV in the presence of 1 μM tetrodotoxin (TTX). Cs-based solution was used, which consisted of (in mM): 40 CsCl, 3.5 KCl, 10 HEPES, 0.05 EGTA, 90 Kgluconate, 1.8 NaCl, 1.7 MgCl2, 2 ATP-magnesium, 0.4 GTP-sodium, and 10 phosphocreatine. The recording batch solution contained (in mM): 140 NaCl, 5 KCl, 10 HEPES, 2 CaCl_2_, 2 MgCl_2_, 10 Glucose, pH 7.4. All cell culture recordings were conducted at room temperature.

### Statistical analysis

Statistical analyses were performed using GraphPad Prism version 10.0 (GraphPad Software, La Jolla, CA). Data are presented as the mean ± standard error of the mean (SEM), unless stated otherwise. The impact of chronic ethanol application was assessed by normalizing the results to the baseline activity in HPRS microglia, with changes in HPRS and LPRS microglia relative to baseline values defined as relative changes. Differences in the effects of various ethanol concentrations in HPRS and LPRS microglia were evaluated using a two-way analysis of variance (ANOVA), followed by post hoc analysis with Fisher’s least significant difference (LSD) test for multiple comparisons. Within the same group of microglia, the effects of ethanol before and after application were compared using unpaired t-tests. Statistical significance was indicated by the following symbols: (*p < 0.05, **p < 0.01, ***p < 0.001), with (NS) denoting no statistically significant difference.

## Acknowledgments

We thank lab members of the Pang laboratory, Dr. Peng Jiang and Dr. Mengmeng Jin for supporting this research. The Child Health Insitute of New Jersey is supported in part by the Robert Wood Johnson Foundation (RWJF grant #74260). We also thank other members of the Pang laboratory and the COGA consortium for their support and valuable comments.

The Collaborative Study on the Genetics of Alcoholism (COGA), Principal Investigators B. Porjesz, V. Hesselbrock, T. Foroud; Scientific Director, A. Agrawal; Translational Director, D. Dick, includes ten different centers: University of Connecticut (V. Hesselbrock); Indiana University (H.J. Edenberg, T. Foroud, Y. Liu, M.H. Plawecki); University of Iowa Carver College of Medicine (S. Kuperman, J. Kramer); SUNY Downstate Health Sciences University (B. Porjesz, J. Meyers, C. Kamarajan, A. Pandey); Washington University in St. Louis (L. Bierut, J. Rice, K. Bucholz, A. Agrawal); University of California at San Diego (M. Schuckit); Rutgers University (J. Tischfield, D. Dick, R. Hart, J. Salvatore); The Children’s Hospital of Philadelphia, University of Pennsylvania (L. Almasy); Icahn School of Medicine at Mount Sinai (A. Goate, P. Slesinger); and Howard University (D. Scott). Other COGA collaborators include: L. Bauer (University of Connecticut); J. Nurnberger Jr., L. Wetherill, X., Xuei, D. Lai, S. O’Connor, (Indiana University); G. Chan (University of Iowa; University of Connecticut); D.B. Chorlian, J. Zhang, P. Barr, S. Kinreich, G. Pandey (SUNY Downstate); N. Mullins (Icahn School of Medicine at Mount Sinai); A. Anokhin, S. Hartz, E. Johnson, V. McCutcheon, S. Saccone (Washington University); J. Moore, F. Aliev, Z. Pang, S. Kuo (Rutgers University); A. Merikangas (The Children’s Hospital of Philadelphia and University of Pennsylvania); H. Chin and A. Parsian are the NIAAA Staff Collaborators. We continue to be inspired by our memories of Henri Begleiter and Theodore Reich, founding PI and Co-PI of COGA, and also owe a debt of gratitude to other past organizers of COGA, including Ting-Kai Li, P. Michael Conneally, Raymond Crowe, and Wendy Reich, for their critical contributions. This national collaborative study is supported by NIH Grant U10AA008401 from the National Institute on Alcohol Abuse and Alcoholism (NIAAA) and the National Institute on Drug Abuse (NIDA).

## Author contributions

ZPP and RPH: conception and design of the research. XL: conducted the experiments, analyzed data, and wrote the manuscript. JL, AK, SZ, JD, and RPH: Analyzed RNA sequence data. AB, SK, ACS, YA, and LC: conducted the experiments and analyzed data. JS, PBB, DL, YL, JLM, CK, WK, AA, PAS, DD, JT, and HJE: sample collection, selection and calculated the AUD-PRS, help with data interpretation. All authors read and approved the manuscript.

## Data and materials availability

RNA sequencing data are available at Gene Expression Omnibus (GEO): GSE255988

