## Supplemental Figures and method for "Polygenic Risk for Alcohol Use Disorder Affects Cellular Responses to Ethanol Exposure in a Human Microglial Cell Model"

Xindi Li *et al.*

**This PDF file includes:**

Supplementary Text
Figs. S1 to S4
Tables S1 to S3
References (95 to 100) (if applicable—these should refer only to references in the SM)

### Supplementary Text

Full details of **bioinformatics analysis of RNA-Seq data**

*Data processing and normalization*

We normalize the sample x gene count matrix using the log (counts per million mapped reads) method. In detail, we denote $r_{ij}$ as our matrix of raw read counts, for samples $j \in \{1, \ldots, 9\}$, and genes $i \in \{1, \ldots, G\}$. For sample $j$ , we calculate its total number of mapped reads:
$R_j = \sum_{i=1}^{G} r_{ij}$.

Next, we calculate the log-counts per million (log-CPM) value for each gene in each sample as:
$y_{ij} = \log_2\left(\frac{r_{ij}+0.5}{R_j+1.0} \times 10^6\right)$.

The normalized data ($y_{ij}$) are then subject to subsequent analyses.

*Differential expression analyses*

We use the *limma+voom method (95,96)* to model the normalized expression level of genes. On the whole, our pipeline consists of three steps: 1) linear modeling (*96*), 2) *voom* variance modeling, and 3) empirical Bayes differential expression analysis.

First, for linear modeling, we consider a nested interaction formula due to the multiple grouping factors in this study: $y \sim 0 + PRS{:}Concentration + PRS{:}CellLine$, where $y_{ij}$ represents the normalized gene expression. Next, we assume that: $E(y_{ij}) = \mu_{ij} = x_j\beta_i$, where $x_j$ represents the vector of grouping factors aforementioned for sample $j$ , and $\beta_i$ represents the coefficient of gene $i$ that represents the log2-fold-changes (log2FC) in expression between the grouping factors.

Second, we estimate the mean-variance relationship of the gene log-counts so that we obtain the precision weights of genes. In detail, we calculate the fitted counts: $E(r_{ij}) = E(\mu_{ij}) + \log_2(R_j + 1) - \log_2(10^6)$, where $E(\mu_{ij})$ stands for fitted log-CPM values, which is calculated using the linear regression coefficient estimates $\beta_i$ and covariates $x_j$. Next, to retain a smooth mean-variance trend, we fit a LOWESS curve (*97*) to the square-root of residual standard deviations obtained from the linear model as a function of mean log-counts across all samples. Therefore, we can calculate the precision weights:

$$\omega_{ij} = lo\left(E(r_{ij})\right)$$

where $lo(\cdot)$ represents the piecewise linear function defined by the LOWESS curve.

Finally, the log-CPM values $y_{ij}$ and their associated weights $w_{ij}$ are then subject to the *limma* standard linear modeling and empirical-Bayes-based differential expression analysis pipeline. To this end, the significant genes were defined as those with an FDR-adjusted *P* value < 0.05 using empirical Bayes F-tests. Therefore, we obtain the lists of genes that are differentially expressed between different grouping factors.

*Gene set enrichment analyses*

We performed the enrichment analysis using pathways in the Gene Ontology and REACTOME Database (MSigDB v2023.1.Hs, C2:CP:REACTOME, C5:GO:BP) (*96)*. The analyses were implemented using the R package ClusterProfiler *(98)*. The enriched pathways were defined as those with an FDR-adjusted $P$ value < 0.05.

*Identifying similarity between bulk RNA-Seq data with single-cell RNA-Seq data of microglia*
To cross-validate the identification of our microglia samples on a transcriptomic level, we compared them with a publicly available single-cell RNA-Seq data set that comprises eight cell types from adult human brain samples, including microglia cells *(99)*. Briefly, we first identified the top 10 marker genes of each cell type following the standard *Seurat* workflow *(100)*. We then calculated the median expression level of these marker genes in each cell type. Finally, we performed the Pearson correlation tests between the median expression level of cell-type-specific marker of the eight cell types and their expression levels in each hiPSC-derived microglial sample.

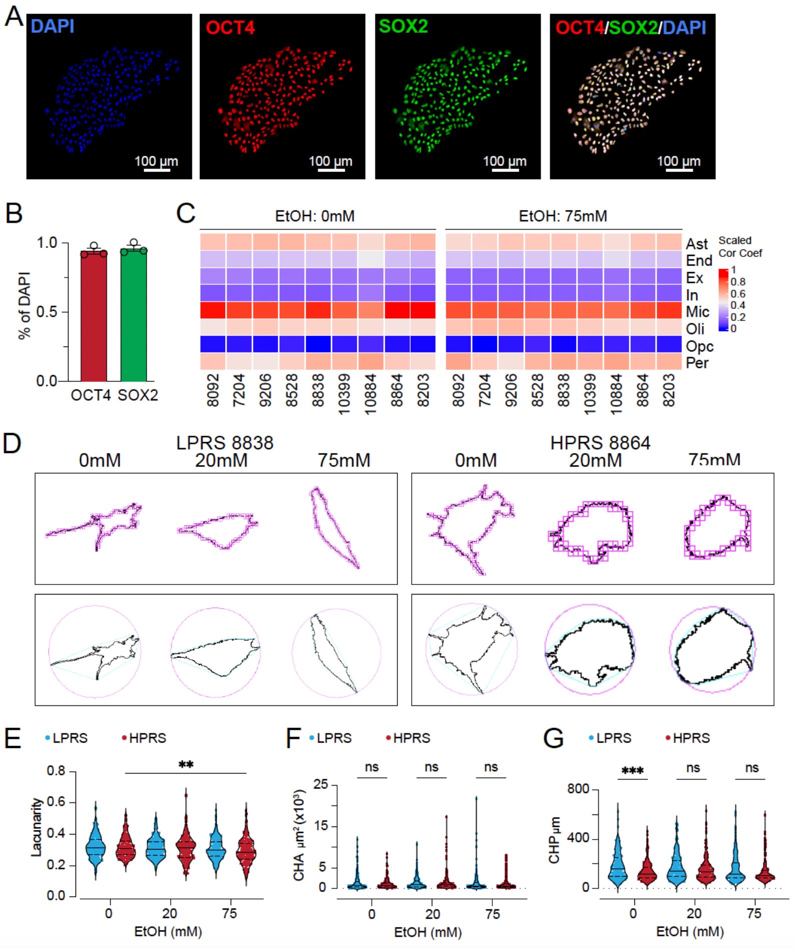

**Fig. S1. Characterization of iPSC derived PMPs and microglia**

**(A)** Representative images of OCT4 and SOX2 expression in iPSCs. **(B)** Quantification of $OCT4^+$ and $SOX2^+$ cells in one pair of high-PRS and low-PRS (HPRS and LPRS) iPSC lines (n = 3). **(C)** The correlation coefficient between the median expression levels of cell-type-specific genes in each cell type of scRNA-Seq data (rows) and their expression levels in each iPSC-derived microglial sample (columns, 5 LPRS lines, 4 HPRS lines). Ast: astrocytes; End: endothelial cells; Ex: excitatory neurons; In: inhibitory neurons, Mic: microglia; Oli: oligodendrocytes; Opc: oligodendrocyte precursor cells; Per: pericytes. **(D)** Upper: illustrating the box-counting method used for fractal dimension and lacunarity calculations. Lower: the associated convex hull (blue) and enclosing circle (pink) for corresponding outline shapes are used to calculate laucunarity, span ratio, and. (E-G) Morphological analysis of microglia binary outlines using FracLac ImageJ. n = 177 cells/HPRS, 211 cells/LPRS for 0 mM; n = 188 cells/HPRS, 214 cells/LPRS for 20 mM; n = 164 cells/HPRS, 219 cells/LPRS for 75 mM. * < 0.05, ** < 0.01, *** < 0.001. Data are presented as medium ± quartiles.

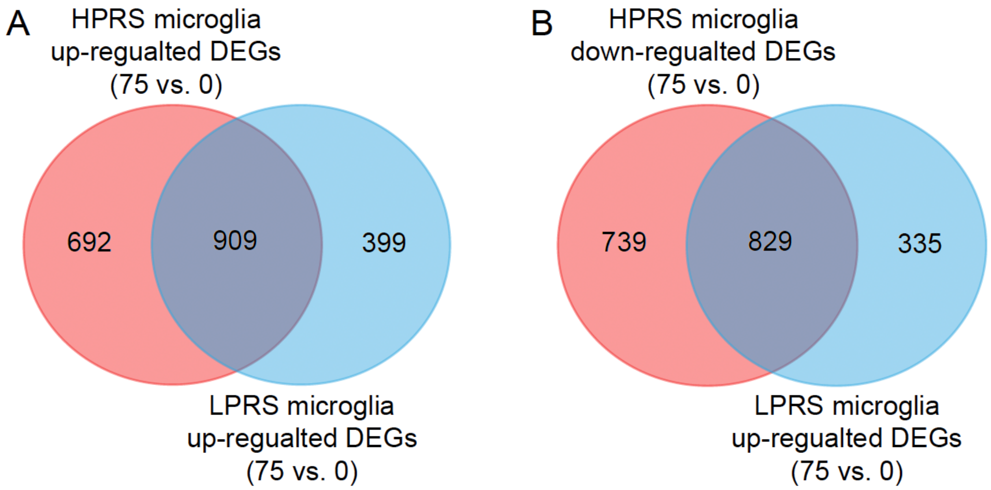


**Fig. S2. Overlapping DEGs between HPRS and LPRS microglial cells**

**(A)** Ven plot illustrating the overlap of upregulated DEGs between HPRS and LPRS microglial cells. **(B)** Ven plot illustrating the overlap of downregulated DEGs between HPRS and LPRS microglial cells.

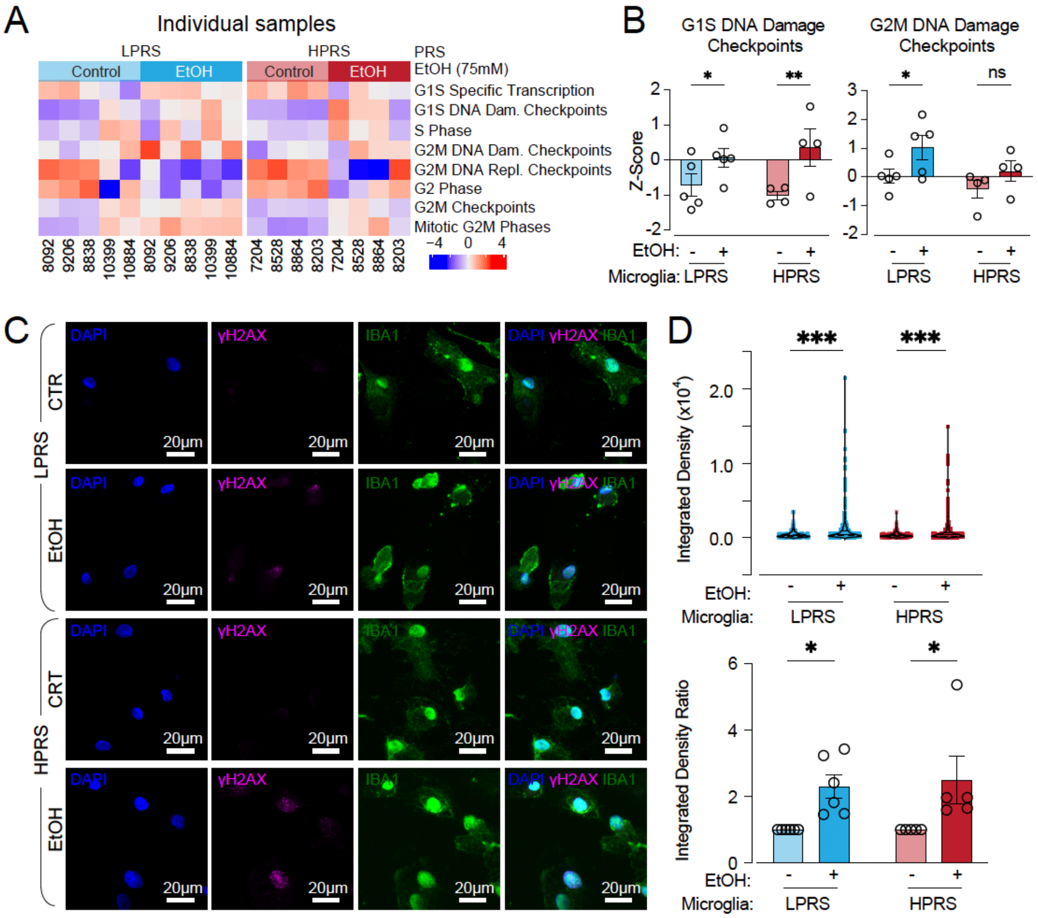


**Fig. S3. DNA damage in HRPS vs. LPRS human microglia after ethanol exposure**

**(A)** Heatmap illustration of the GSEA terms related to the cell cycle in both HPRS (red, n = 4) and LPRS (blue, n = 5) microglia, both before and after ethanol exposure. **(B)** Box plots illustrating the differential expression of GSEA terms related to the cell cycle in microglia from both HPRS (red, n = 4) and LPRS (blue, n =5) lines before and after ethanol treatment. LPRS/HPRS lines = 6/5. * < 0.05, ** < 0.01. Data are presented as mean ± SEM. **(C)** Representative images of γ-H2AX$^{+}$ and IBA1$^{+}$ microglia derived from PRS lines following treatment with ethanol (0 mM and 75 mM). **(D)** Upper: the quantification of γ-H2AX$^{+}$ fluorescence integrated density. n = 357 cells/5 HPRS lines and 421 cells/6 LPRS lines for 0 mM; n = 392 cells/5 HPRS lines, 447 cells/6 LPRS lines for 75 mM; ** < 0.01, *** < 0.001. Data are presented as mean ± SEM. Lower: the quantification of γ-H2AX$^{+}$ corrected fluorescence integrated density normalized to each control (0 mM) replicate. ** < 0.01, *** < 0.001. Data are presented as mean ± SEM.

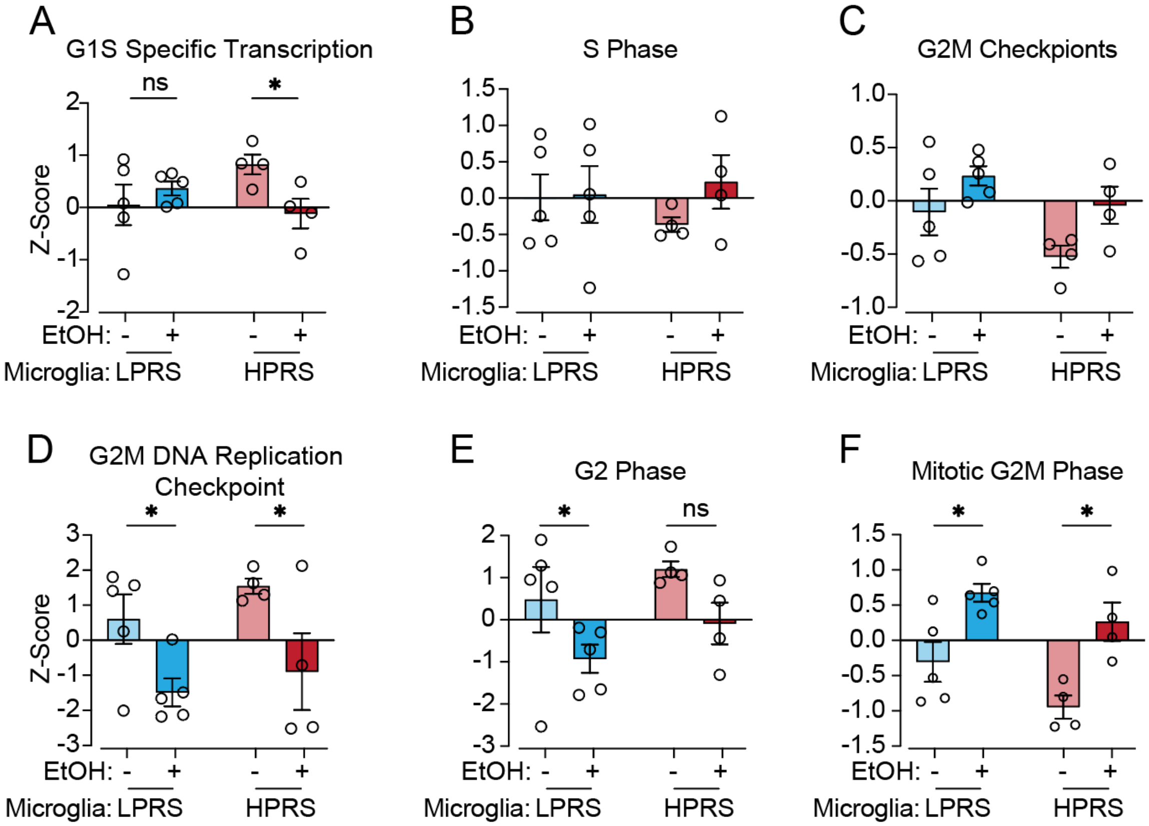


**Fig. S4. Expression of GSEA terms related to the cell cycle in HPRS and LPRS microglial cells after ethanol exposure**

**(A-F)** Box plots illustrating the differential expression of GSEA terms related to the cell cycle in microglial cells from both HPRS (red, n=4 lines) and LPRS (blue, n=5 lines) lines before and after Ethanol treatment. * < 0.05, ** < 0.01. Data are presented as mean ± SEM.

**Table S1. Clinical characteristics of individuals enrolled for iPSC generation.**

| | Gender | Age | ID | PRS %ile | AUD | Bulk seq | Morphology & ICC | Phagocytosis assay | iN & Microglia Co-culture |
|---|---|---|---|---|---|---|---|---|---|
| HPRS | male | 38 | 8864 | 97.60 | Yes | Yes | Yes | Yes | No |
| HPRS | male | 38 | 8528 | 97.70 | Yes | Yes | Yes | Yes | No |
| HPRS | male | 38 | 8203 | 98.70 | Yes | Yes | Yes | Yes | Yes |
| HPRS | female | 31 | 7204 | 98.80 | Yes | Yes | Yes | Yes | Yes |
| HPRS | female | 39 | 8260 | 99.40 | Yes | No | Yes | Yes | Yes |
| HPRS | male | 56 | 8059 | 95.56 | Yes | No | No | Yes | Yes |
| HPRS | female | 28 | 8098 | 99.76 | Yes | No | No | Yes | Yes |
| HPRS | male | 32 | 10179 | 75.50 | Yes | No | No | Yes | Yes |
| LPRS | male | 27 | 0884 | 1.40 | No | Yes | Yes | Yes | No |
| LPRS | male | 29 | 8838 | 2.80 | No | Yes | Yes | Yes | No |
| LPRS | female | 22 | 9206 | 0.90 | No | Yes | Yes | Yes | No |
| LPRS | female | 33 | 8092 | 5.10 | No | Yes | Yes | Yes | Yes |
| LPRS | female | 38 | 0399 | 0.89 | No | Yes | Yes | Yes | Yes |
| LPRS | male | 27 | 9618 | 0.40 | No | No | Yes | Yes | Yes |
| LPRS | male | 26 | 0028 | 13.09 | No | No | No | Yes | Yes |
| LPRS | female | 22 | 0196 | 4.24 | No | No | No | Yes | Yes |
| LPRS | female | 33 | 0929 | 1.44 | No | No | No | Yes | Yes |
| LPRS | male | 28 | 9618 | 0.40 | No | No | No | Yes | Yes |

Note: HPRS: High polygenic risk scores; LPRS: Low polygenic risk scores; ICC: Immunological cytochemistry; Age: Subject age at time of lymphocyte collection.

### Table S2. Gene expression count of ADH and ALDH genes in Bulk sequence from iPSC-derived microglia

| Gene ID | Gene name | 8092 | | 7204 | | 9206 | | 8528 | | 8838 | | 10399 | | 10884 | | 8864 | | 8203 | |
|---|---|---|---|---|---|---|---|---|---|---|---|---|---|---|---|---|---|---|---|
| | | 0mM | 75mM | 0mM | 75mM | 0mM | 75mM | 0mM | 75mM | 0mM | 75mM | 0mM | 75mM | 0mM | 75mM | 0mM | 75mM | 0mM | 75mM |
| ENSG00000187758 | ADH1A | 0 | 0 | 0 | 0 | 0 | 0 | 4 | 6 | 0 | 0 | 0 | 0 | 0 | 0 | 0 | 0 | 0 | 0 |
| ENSG00000197894 | ADH5 | 1618 | 1859 | 2084 | 1890 | 1612 | 1645 | 1572 | 1540 | 1579 | 1584 | 1522 | 1946 | 1686 | 1329 | 1461 | 1443 | 1424 | 1150 |
| ENSG00000147576 | ADHFE1 | 15 | 44 | 29 | 25 | 21 | 23 | 8 | 23 | 20 | 29 | 27 | 28 | 18 | 19 | 19 | 24 | 27 | 8 |
| ENSG00000161618 | ALDH16A1 | 1148 | 950 | 3112 | 1838 | 1760 | 1424 | 1812 | 1486 | 1088 | 1039 | 1929 | 1870 | 2189 | 1803 | 1613 | 1526 | 1754 | 1487 |
| ENSG00000059573 | ALDH18A1 | 159 | 140 | 177 | 188 | 131 | 142 | 197 | 185 | 180 | 159 | 190 | 283 | 168 | 253 | 200 | 210 | 197 | 149 |
| ENSG00000165092 | ALDH1A1 | 3 | 1 | 43 | 15 | 79 | 19 | 224 | 69 | 71 | 50 | 308 | 126 | 445 | 244 | 330 | 172 | 47 | 22 |
| ENSG00000128918 | ALDH1A2 | 7704 | 1993 | 8015 | 2556 | 3436 | 1006 | 10190 | 2009 | 2913 | 723 | 3997 | 2062 | 694 | 224 | 3563 | 1903 | 9678 | 4732 |
| ENSG00000137124 | ALDH1B1 | 63 | 27 | 188 | 103 | 137 | 90 | 91 | 40 | 75 | 42 | 35 | 58 | 151 | 108 | 52 | 64 | 171 | 68 |
| ENSG00000136010 | ALDH1L2 | 74 | 93 | 91 | 161 | 51 | 67 | 86 | 129 | 119 | 113 | 66 | 90 | 173 | 315 | 123 | 111 | 48 | 73 |
| ENSG00000111275 | ALDH2 | 657 | 848 | 697 | 952 | 692 | 802 | 857 | 1016 | 930 | 977 | 1179 | 1683 | 837 | 810 | 904 | 1558 | 310 | 538 |
| ENSG00000072210 | ALDH3A2 | 634 | 620 | 903 | 848 | 826 | 804 | 770 | 701 | 827 | 685 | 771 | 985 | 740 | 660 | 867 | 828 | 679 | 515 |

| | | | | | | | | | | | | | | | | | | | |
|---|---|---|---|---|---|---|---|---|---|---|---|---|---|---|---|---|---|---|---|
| ENSG00000006534 | ALDH3B1 | 1688 | 1745 | 2464 | 1745 | 1561 | 1515 | 1711 | 1713 | 1602 | 1461 | 1714 | 1844 | 2117 | 1993 | 1756 | 1771 | 2037 | 1401 |
| ENSG00000159423 | ALDH4A1 | 318 | 273 | 639 | 435 | 581 | 387 | 520 | 407 | 360 | 347 | 392 | 327 | 523 | 381 | 427 | 415 | 330 | 295 |
| ENSG00000112294 | ALDH5A1 | 188 | 95 | 449 | 206 | 229 | 142 | 298 | 82 | 212 | 83 | 376 | 271 | 54 | 95 | 289 | 192 | 742 | 204 |
| ENSG00000119711 | ALDH6A1 | 201 | 226 | 359 | 384 | 307 | 243 | 298 | 309 | 300 | 279 | 475 | 444 | 465 | 366 | 202 | 182 | 348 | 279 |
| ENSG00000164904 | ALDH7A1 | 463 | 543 | 719 | 711 | 709 | 678 | 513 | 737 | 623 | 586 | 593 | 807 | 623 | 708 | 611 | 770 | 489 | 521 |
| ENSG00000143149 | ALDH9A1 | 1599 | 1478 | 2154 | 1620 | 1820 | 1711 | 1849 | 1518 | 1437 | 1341 | 2162 | 2219 | 1703 | 1510 | 1432 | 1401 | 1657 | 1354 |
| ENSG00000180011 | ZADH2 | 993 | 855 | 1671 | 1392 | 1374 | 1249 | 1345 | 1158 | 1063 | 979 | 1343 | 1571 | 709 | 601 | 1149 | 992 | 1326 | 816 |

**Table S3. General information of sequencing samples**

| sample | library | raw_ reads | raw_ bases | clean_ reads | clean_ bases | error _rate | Q20 | Q30 | GC_pct |
|---|---|---|---|---|---|---|---|---|---|
| LPRS groups | | | | | | | | | |
| 8092 (0mM) | CRAS230023998-1r | 42818486 | 6.42G | 41406848 | 6.21G | 0.02 | 97.99 | 94.45 | 52.2 |
| 8092 (75mM) | CRAS230024000-1r | 42539472 | 6.38G | 41198130 | 6.18G | 0.02 | 98.06 | 94.54 | 52.51 |
| 8838 (0mM) | CRAS230024010-1r | 45456668 | 6.82G | 44053674 | 6.61G | 0.02 | 98.11 | 94.67 | 52.46 |
| 8838 (75mM) | CRAS230024012-1r | 43116898 | 6.47G | 41864472 | 6.28G | 0.02 | 97.97 | 94.35 | 53.01 |
| 10399 (0mM) | CRAS230024013-1r | 48919870 | 7.34G | 47371032 | 7.11G | 0.02 | 98.06 | 94.53 | 52.5 |
| 10399 (75mM) | CRRA230024015-1a | 64576460 | 9.69G | 63074980 | 9.46G | 0.03 | 97.56 | 93.78 | 54.05 |
| 10884 (0mM) | CRRA230024016-1a | 51922120 | 7.79G | 50675938 | 7.6G | 0.03 | 97.7 | 93.91 | 53.13 |
| 10884 (75mM) | CRRA230024018-1a | 46247992 | 6.94G | 45096246 | 6.76G | 0.03 | 97.59 | 93.65 | 53.62 |
| 9206 (0mM) | CRAS230024004-1r | 46855900 | 7.03G | 45152358 | 6.77G | 0.02 | 98.13 | 94.75 | 52.59 |
| 9206 (75mM) | CRAS230024006-1r | 45254386 | 6.79G | 43866134 | 6.58G | 0.02 | 98.08 | 94.6 | 52.92 |
| HPRS groups | | | | | | | | | |
| 8528 (0mM) | CRAS230024007-1r | 50713000 | 7.61G | 49252646 | 7.39G | 0.02 | 97.93 | 94.28 | 52.68 |
| 8528 (75mM) | CRAS230024009-1r | 50651220 | 7.6G | 48944528 | 7.34G | 0.03 | 97.92 | 94.27 | 53.17 |
| 8864 (0mM) | CRRA230024019-1a | 44768638 | 6.72G | 43974814 | 6.6G | 0.03 | 97.57 | 93.6 | 52.9 |
| 8864 (75mM) | CRRA230024021-1a | 51643684 | 7.75G | 50863276 | 7.63G | 0.03 | 97.74 | 93.94 | 53.24 |
| 8203 (0mM) | CRRA230024022-1a | 49572568 | 7.44G | 48741338 | 7.31G | 0.03 | 97.67 | 93.9 | 52.86 |
| 8203 (75mM) | CRRA230024024-1a | 42315166 | 6.35G | 41459172 | 6.22G | 0.03 | 97.78 | 94.06 | 53 |
| 7204 (0mM) | CRAS230024001-1r | 65632920 | 9.84G | 61952374 | 9.29G | 0.02 | 98.07 | 94.45 | 54.21 |
| 7204 (75mM) | CRAS230024003-1r | 54570298 | 8.19G | 52910082 | 7.94G | 0.03 | 97.9 | 94.16 | 52.61 |
